# A Hybrid Stochastic Model of the Budding Yeast Cell Cycle

**DOI:** 10.1101/709691

**Authors:** Mansooreh Ahmadian, John J. Tyson, Jean Peccoud, Yang Cao

**Affiliations:** Department of Computer Science, Virginia Tech, Blacksburg, VA, USA; Department of Biological Sciences, Virginia Tech, Blacksburg, VA, USA; GenoFAB Inc, Fort Collins, CO, USA

**Author notes:** Correspondence: Yang Cao and John J. Tyson.

## Abstract

The growth and division of eukaryotic cells are regulated by complex, multi-scale networks. In this process, the mechanism controlling cell cycle progression has to be robust against inherent noise in the system. In this paper, a hybrid stochastic model is developed to study the effects of noise on the control mechanism of the budding yeast cell cycle. The modeling approach leverages, in a single multi-scale model, the advantages of two regimes: 1) the computational efficiency of a deterministic approach, and 2) the accuracy of stochastic simulations. Our results show that this hybrid stochastic model achieves high computational efficiency while generating simulation results that match very well with published experimental measurements.

## 1 Introduction

The eukaryotic cell cycle is a complex process by which a growing cell replicates its DNA and divides into two cells, each capable of repeating the process. Progression through the cycle is controlled by networks of genes, mRNAs, and proteins, with interactions that can be modeled as chemical reaction channels. To unravel the complex dynamics of multi-scale reaction networks in higher organisms such as human cells, it is advisable to study single-cell organisms with molecular regulatory networks that are similar yet simpler. For instance, experimental studies and mathematical models of frog eggs [1, 2], fission yeast [3, 4], and budding yeast [5, 6] have shed light on mechanisms of cell cycle regulation in the cells of higher organisms. Extensive experimental studies have been conducted particularly on the budding yeast (Saccharomyces cerevisiae) to explore gene regulation and signaling pathways of relevance to cell growth and division [7–9]. Moreover, various modeling approaches, such as deterministic models [10–12], Boolean networks [13–19], and stochastic models [20–27], have been adopted to explore the roles of different gene and protein interactions in robust progression through the cell cycle.

Among these models, a deterministic approach is most common. In this approach, the time-dependent variation of each molecular species in the biochemical reaction network is described by a nonlinear ordinary differential equation (ODE), in which the concentration of the substance is considered as a continuous quantity that evolves deterministically over time. However, the time-evolution of molecular species within the confined volume of a budding yeast cell (about 30 fL at birth) is not deterministic. Therefore, in spite of being able to reproduce certain average characteristics of cell cycle progression in yeast cell populations, a continuous-deterministic model cannot reproduce the cell-to-cell variability observed in wet-lab experiments [28, 29]. For instance, Di Talia *et al.* [29] have reported that the coefficient of variation 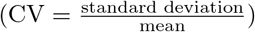 for G1 time of bundding yeast cells (growing on glucose) is 50%.

To capture such high levels of variability, stochastic models have been built using different strategies to incorporate intrinsic and extrinsic sources of noise. In an early stochastic model of the fission yeast cell cycle proposed by Sveiczer *et al.* [30], extrinsic noise was introduced by assuming some sloppiness in the partitioning of cell volume and nucleus volume to daughter cells at division. A later model by Steuer [31] examined the roles of intrinsic noise in cell cycle progression by adding Gaussian noise to reaction rate equations in a deterministic model. These approaches, however, do not adequately explain the root source of cell-cycle variability in yeast cells, which lies in molecular fluctuations at the level of gene expression [32–34]. To capture such molecular-level noise, more accurate stochastic methods are required to explicitly model fluctuations in molecular interactions. For this purpose, the best method to implement fluctuating molecular interactions is the *stochastic simulation algorithm* (SSA) proposed by Gillespie [35]. Gillespie’s algorithm is a Monte-Carlo approach that numerically simulates the temporal firing of every single reaction in a chemical reaction network. An assumption of Gillespie’s method is that the propensity of every reaction in the model is described by mass-action kinetics. This becomes an issue for us because most deterministic models of cell cycle regulation, such as those presented in [5, 11, 12, 36], incorporate complex rate laws including Michaelis-Menten kinetics, Hill functions, and ultra-sensitive switches. These complex phenomenological rate laws are used in deterministic models to provide sufficient nonlinearity in reaction kinetics to create bistable switches that flip on and off during progression through the cell cycle. Converting a deterministic model into a stochastic model suitable for Gillespie’s SSA by ‘unpacking’ complex rate laws into elementary reactions is a difficult problem fraught with uncertainties [37].

To address this challenge, several approaches have been tried. The simplest approach, used for example by Mura & Csikasz-Nagy [38], treats all complex rate laws directly as propensity functions of reactions and then applies the SSA. This approximation is subject to considerable errors [37, 39]. For example, Ball *et al.* [21] found that the variability they observed in wet-lab measurements could not be generated by this greatly simplified stochastic approach, unless some unrealistic parameter values were chosen. Later Kar *et al.* [20] tried to unpack Michaelis-Menten rate laws in a small (three-variable), deterministic model of the budding yeast cell cycle [5]. Unpacking resulted in a much more complicated system with 19 species and 47 reactions. Although this simple model (with only a few key cell-cycle genes) could generate noise levels that match wet-lab measurements for a few key characteristics of the cell cycle, it is not feasible (in our experience) to apply this approach to more complex models with substantially more genes and proteins. Instead, we have pursued an approach in which the molecular controls of the budding yeast cell cycle are modeled directly in terms of elementary reactions (governed by the law of mass-action) [25, 26]. A great advantage of this approach is that the newly designed deterministic model can be converted into its corresponding stochastic version without any approximation. A disadvantage of this approach is that we cannot re-purpose our original deterministic models, which had been carefully designed and parametrized to explain a broad scope of experimental observations. Furthermore, to model the phosphorylation and dephosphorylation reactions that play such important roles in cell cycle progression introduces substantial complexity into the system. Recently we have considered a new approach that sidesteps the complications of elementary reactions and mass-action rate laws and that employs a Langevin-type simulation of noisy gene expression [23]. This approach, though promising, also requires an overhaul of the original deterministic models. In order to take advantage of existing deterministic models in a framework that permits accurate stochastic simulations without ‘unpacking’, we explore a particular hybrid approach in this paper.

Gillespie’s SSA simulates every single reaction firing. In general, the time complexity of this algorithm scales proportionally with the number of reaction firings. Consequently, SSA-based models involve substantial computational complexity if a reaction network involves many fast reactions. To reduce the high computational cost of the SSA, many optimization methods [40–44] and approximation methods [45–49] have been proposed. Among them the *hybrid stochastic* approach, originally proposed by Haseltine and Rawling (HR) [47], performs well because it takes advantage of the multi-scale features common in biochemical reaction networks. The main idea of the HR hybrid approach is to partition the dynamical system into *fast* and *slow* reactions, based on the relative time scale of each reaction and the abundances of the reactants. Fast reactions, which fire frequently and often involve high-abundance species, are partitioned into the deterministic (ODE) regime. Meanwhile, slow reactions, which are often found at the gene-expression level, fire much less frequently and are therefore simulated using the SSA. This approach was first applied by Liu *et al.* [22] on the simple three-variable model of the budding yeast cell cycle, originally studied by Kar *et al.* [20]. By partitioning all gene-expression reactions into the slow (SSA) regime and all protein-level dynamics into the fast (ODE) regime, Liu *et al.* [22] were able to reproduce the noise levels that Kar *et al.* [20] achieved by unpacking the original system into a much more complex one. This success motivated us to apply this approach to the very comprehensive, accurate, and complex deterministic model of Chen *et al.* [11].

Simulation results demonstrate that, while achieving high computational efficiency, our hybrid model still matches up well with experimental measurements of the variability of cell-cycle-related properties (cycle time, cell size, correlation coefficients), protein and mRNA abundances, and phenotypes of more than 100 mutant strains of budding yeast. Moreover, our simulations shed light on the ‘partial’ viability of mutant strains such as *CLB*2*db*Δ *clb*5Δ.

## 2 Model

### 2.1 Deterministic model

A comprehensive continuous-deterministic model of the budding yeast cell cycle was developed by Chen *et al.* [11] in 2004. By integrating the findings of decades of experimental studies the Chen model provides an accurate mathematical description of the cell division cycle of budding yeast. The protein regulatory network of Chen’s model focuses primarily on the mutual antagonism between mitotic B-type cyclins (Clb1-6) and G1 phase stabilizers (Cdh1, Sic1, and Cdc6). During the growth and division of yeast cells, this antagonism leads to transitions between two coexisting steady states called START (G1→ S) and EXIT (M→ G1). A detailed description of the budding yeast cell cycle model is given in Supplementary Information.

Chen’s mathematical model reproduces the average cell-cycle properties (including cycle time, G1 duration, and cell size at division) of wild-type budding yeast cells and the variant cell-cycle phenotypes of more than 100 mutant strains. Our goal is to develop a hybrid (stochastic-deterministic) version of this large regulatory network, in order to quantify the variabilities observed in cell cycle characteristics and mutant phenotypes within a computationally efficient framework.

Since Chen’s model is formulated in terms of normalized (dimensionless) concentrations of proteins, the first step to this goal is to convert the state variables of Chen’s model into integer numbers of molecules per cell. This conversion facilitates comparison of our numerical simulation results with observed data from single-cell experiments. Furthermore, it is necessary because, in Gillespie’s SSA, state variables are discrete (species populations) rather than continuous (species concentrations). Since a hybrid model involves both SSA and ODEs, it is important that we assure consistency between units of state variables in both the stochastic and deterministic regimes. Therefore, we calculate *S_i_*, the number of molecules of species *i* in a cell, from the corresponding normalized concentration, [S_*i*_]_n_, by (1):

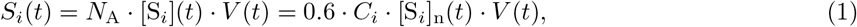

where [S_*i*_] is the actual concentration of species *i* (in nanomoles/liter = 10^−9^ mol/L), *C_i_* is the ‘characteristic’ concentration of species *i* (used to convert between actual concentration and ‘normalized’ concentration), *V* (*t*) is the volume of the cell (in femtoliters = 10^−15^ L), and *N*_A_ = 0.6 is Avogadro’s number (when concentration is expressed in nM and volume in fL). One simplifying assumption made in published models [21, 23] is to use a constant volume for the size of cell. However, this unrealistic assumption introduces errors into the model because cell size (*V*) increases exponentially during a cycle 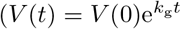, where *k_g_* is the specific growth rate of yeast cells).

Second, we extend the protein regulatory network in Chen’s model to include the dynamics of 11 regulated and 8 unregulated mRNAs. This extension is necessary because the major source of intrinsic noise in yeast cells is the small number of mRNA molecules per cell per gene [28]. Experimental observations [29] in yeast cells with increased dosage of genes suggest that the dominant source of variability with respect to cell-cycle time and cell size at division is the low copy number of mRNA and protein molecules in a cell, specifically in G1 phase. However, Chen’s model did not incorporate the turnover of mRNA molecules, and thus it cannot account for fluctuations stemming from transcriptional noise. For these reasons, Chen’s original model must be supplemented with appropriate synthesis and degradation rates for each mRNA, as well as realistic rates of translation from mRNA to protein. In our model, based on experimental observations in [50], we assigned half-life times for mRNAs in the range of 5-10 min, except for *mCln*2 and *mClb*2, which were assigned shorter half-lives (3 and 2 min, respectively). The synthesis rate of each mRNA was then estimated to match the mRNA average-abundance measurements in Ball *et al.* [28].

Third, we modified Chen’s model by introducing ODEs for the concentrations of Cln3 and Bck2 proteins. In Chen’s original model, the normalized concentration of Cln3 and Bck2 were assumed to be given by steady-state algebraic equations (2) and (3),

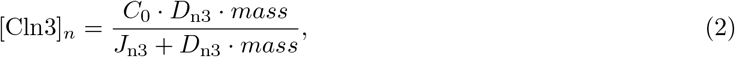

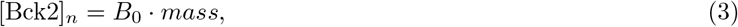

where *C*_0_ determines the maximum concentration of Cln3, *D*_n3_ is the dosage of the *CLN* 3 gene, *J*_n3_ and *B*_0_ are constants, and *mass* is the ‘size’ of a cell. We replaced the algebraic equations (2) and (3) by ODEs in (4) and (5).

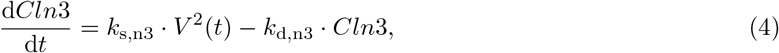

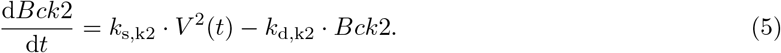

The synthesis (*k*_s,n3_, *k*_s,k2_) and degradation (*k*_d,n3_, *k*_d,k2_) rate constants were estimated so that the half-lives and average abundances of these proteins match with experimental data [51]. The reason for this change is to model the unbalanced partitioning of Cln3 molecules between daughter and mother cells at cell division. According to experimental observations, the concentration of Cln3 in a new-born daughter cell is about 3 times less than its concentration in the mother cell [52, 53], indicating that mother cells get more than their ‘fair share’ of Cln3 molecules at cell separation. As a consequence of this unequal partitioning of Cln3 between mother and daughter cells at division, the G1 time of mother cells is much shorter and the G1 time of daughter cells is much longer (on average) than would otherwise be expected. By including Cln3 and Bck2 as state variables in the model, we can apply an asymmetric partitioning rule with ratio of 20:80 to daughter and mother cells at cell division. We note that this ratio is set to 40:60 for all other proteins and mRNAs, according to observations in [29].

Finally, we comment that the quadratic dependence of Cln3 and Bck2 synthesis rates on cell size is introduced to account for the major influence that these two proteins have on cell size at the G1/S transition [54, 55]. Because the rate of synthesis of these two proteins increases quadratically with cell volume, there is a strong size control on the G1/S transition in our model.

In summary, the variables, equations, parameter values and reaction propensities in our model are provided in Supplementary Tables S1-S4 in Supplementary Text.

### 2.2 Hybrid Stochastic Model

As we mentioned in Section 1, the regulatory network of the budding yeast cell cycle is a multi-scale system: both the numbers of molecules of mRNAs and proteins and the propensities of individual reactions vary by orders of magnitude. For instance, in budding yeast cells, there are 500-5,000 copies of each protein encoded by only 5-10 copies of the corresponding mRNA. Furthermore, the synthesis and degradation of mRNA species occur much less frequently than the phosphorylation and dephosphorylation of proteins in the cell. The HR hybrid method leverages these large scale differences to improve the efficiency of stochastic simulations without sacrificing accuracy of the computations. The HR method divides the system into subsystems, each including species and reactions with similar scales, and applies an appropriate simulation method to each subsystem. This partitioning is done by using predefined thresholds for propensities of reactions and abundances of reactants. In this way the system is divided into four disjoint regions: (I) slow reactions with low-abundance reactants, (II) slow reactions with high-abundance reactants, (III) fast reactions with low-abundance reactants, and (IV) fast reactions with high-abundance reactants. Then an appropriate simulation method is chosen for each region [22, 41, 47]. We follow the strategy proposed Liu *et al.* [22] where the dynamics of all mRNAs (region I) is simulated by SSA, and the other three regions (II, III, and IV) are modeled with ODEs. We shall refer to this partitioning as the ‘Liu strategy’.

**Algorithm 1.**
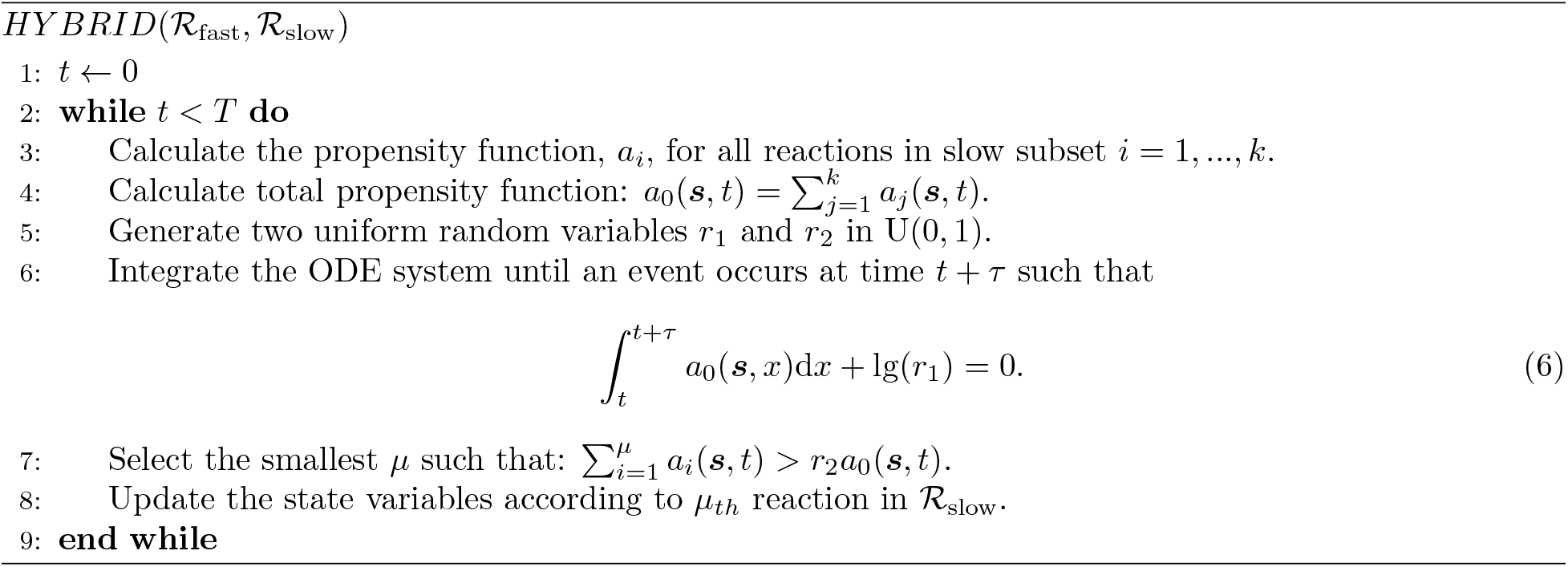
Hybrid Stochastic Simulation Algorithm.

#### Hybrid Stochastic Simulation Algorithm

Consider a well-stirred system with *N* species in a set 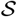 that interact with each other through *M* reaction channels in a set 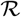. The reactions in 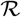 are partitioned into two disjoint subsets of fast and slow reactions denoted by 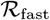 and 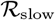, respectively. The subset 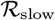 includes *k* reactions which are simulated using SSA, while the remaining *M*-*k* fast reactions in 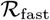 are governed by ODEs. Let *a_j_*(***s***, *t*) be the propensity function of the *j*-th reaction in 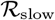, where ***s*** = (*S*_1_(*t*), …, *S*_*N*_ (*t*)) is the state vector with each element *S_i_*(*t*) representing the number of molecules of species *i* at time *t*. In addition, let *v_j_* = (*v*_*j*1_, …, *v_jN_*) be the state-change vector of the *j*-th reaction, where *v_ji_* denotes the change in the population of species *i* when reaction *j* fires. Let *τ* be the jump interval to the next slow reaction and *μ* be the index of the reaction that fires. The algorithm only needs to simulate the firings of slow reactions, while integrating the fast subset of ODEs simultaneously in (6). When a slow reaction fires, the corresponding state variables are updated. In this way the hybrid algorithm generates trajectories of state variables as the system proceeds in time. More details on implementation can be found in [56–59].

In order to demonstrate the scale difference in our partitioning strategy, we approximate the propensity function of every reaction by its corresponding rate law function (obtaining a stochastic model with 145 reactions) and track the firing frequency of each of these reactions in a test run of Gillespie’s SSA. Of 18 million reactions fired in one cell cycle, only about 34,000 (0.2%) involve mRNA turnover, and 99.8% represent fast reactions of protein post-translational modifications. Based on this test run, we estimate that our HR hybrid scheme will run at least 100 times faster than a brute-force Gillespie simulation of a fully stochastic model. In section 3 we show that our hybrid stochastic model, using the Liu strategy, still generates accurate results that agree well with experimental observations.

**Algorithm 1** proposed in Liu et al. [22] describes the hybrid ODE/SSA algorithm adopted in this paper, which is a variant of the original HR hybrid method [47]. Our hybrid stochastic simulation code that implements algorithm 1 (a FORTRAN file) is available in Supplementary Code.

## 3 Results

Our hybrid stochastic model of the budding yeast cell cycle, consisting of 45 proteins and 19 mRNAs, is simulated by Algorithm 1 to generate sufficiently large populations of mother and daughter cells to estimate the statistical distributions of various cell-cycle-related properties of wild-type cells as well as 122 mutant strains of yeast. We evaluate our model by comparing numerical simulation results with experimental observations from the published literature.

### 3.1 Wild-type cell

Figure 1 A and B show a deterministic simulation of several protein and mRNA species, respectively, in our model of wild-type budding yeast cells. In early G1 phase, once the cell grows to a critical size, Cln3 and Bck2 initiate the START event, i.e., the activation of transcription factors for Cln2 and Clb5 production. Cln2 is the cyclin responsible for bud formation. In addition, Cln2 phosphorylates Sic1 and Cdc6, a pair of cyclin-dependent kinase inhibitors (CKIs). Consequently, Clb5-dependent kinase activity rises and initiates DNA replication in S phase. As CKIs are removed, Clb2 level rises, because Clb2 activates its own transcription factor, Mcm1, in an autocatalytic fashion. Clb2-dependent kinase activity turns off the transcription factors for Cln2 and Clb5 production and is responsible for driving the cell into mitosis (M phase). Clb2 level remains high until metaphase, when the proper attachment of chromosomes to the mitotic spindle activates Cdc20. Cdc20 promotes anaphase (the separation of the two strands of replicated chromosomes to opposite poles of the mitotic spindle). At the same time, Cdc20 promotes degradation of Clb2 and Clb5 and activation of a phosphatase, Cdc14. Cdc14 plays a major role (in budding yeast) in re-establishing the dominance of CKIs in G1 phase, and in replacing Cdc20 by Cdh1 (the protein responsible for Clb2 and Clb5 degradation in G1 phase).

**Figure 1:**
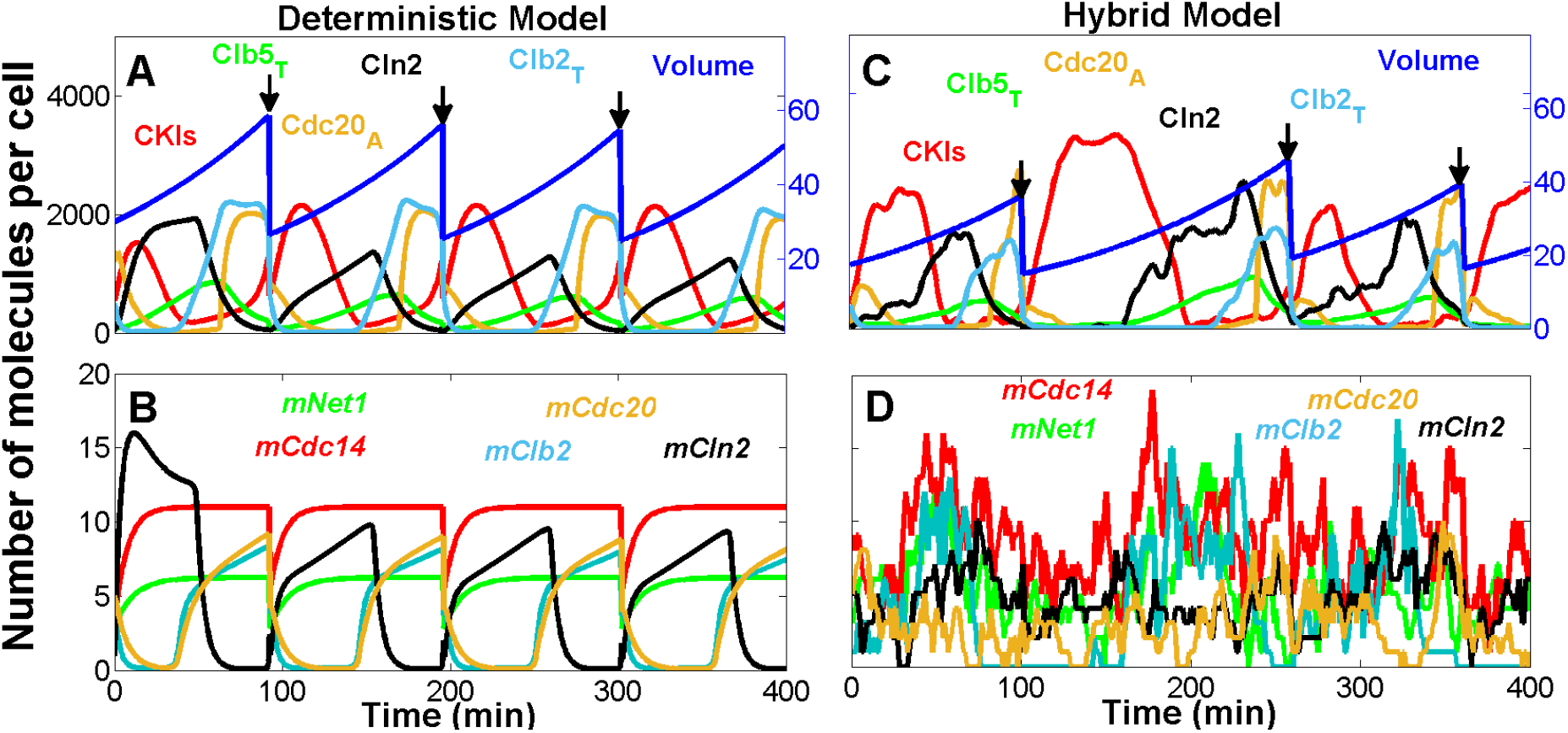
Deterministic and hybrid stochastic simulations of the model. **A and B:** The temporal dynamics of representative proteins (A) and mRNAs (B) generated by the deterministic model. The volume of the cell increases exponentially and is divided (at the arrows) asymmetrically between mother (55%) and daughter cell (45%). **C and D**: Stochastic simulation of the same proteins (C) and mRNAs (D) as in panels A and B, generated by a representative run of the hybrid stochastic model. Similar to the deterministic model, the cell grows exponentially; however, at the time of division all species in the cell, except for Cln3 and Bck2, are partitioned between daughter and mother cells with a 40:60 ratio, according to observations by Di Talia *et al.* [29]. Cln3 and Bck2, which are preferentially retained in mother cells [52, 53], are partitioned with a ratio 20:80 between daughter and mother cells. The daughter cell is tracked from division to division in the this simulation.

Figure 1 C and D show the corresponding stochastic trajectories of our hybrid stochastic model. The stochastic trajectories in panel C correctly simulate the sequence of events predicted by the deterministic trajectories in panel A. Panel D shows considerable fluctuations in the numbers of molecules per cell of five mRNA species: three of which (*mClb*2, *mCdc*20, and *mCln*2) undergo periodic transcription during the cell cycle, and two of which (*mCdc*14 and *mNet*1) are synthesized continuously throughout the cell cycle. Such high variability is expected, due to the stocastic nature of gene expression and the low abundances of mRNA molecules per cell.

We used our hybrid stochastic model to generate more than 10,000 asynchronous wild-type cells growing in glucose medium (mass doubling time about 100 min). This large collection of simulated cells is then used to estimate the distributions of important characteristics of the budding yeast cell cycle, including inter-division time, duration of unbudded phase (G1), duration of budded phase (S-G2-M), and size at birth. Table 1 compares the computed summary statistics of these cell cycle-related properties with experimental data reported by Di Talia *et al.* [29]. The results in Table 1 show that the model accurately reproduces the mean and coefficient of variation of these important properties of the wild-type budding yeast cell cycle.

**Table 1:** Mean and coefficient of variation (CV) for cell cycle properties. Summary statistics including mean and CV computed from a simulation of the hybrid stochastic model are compared with experimental observations reported by Di Talia *et al.* [29]. *T*_div_, *T*_G1_, *T*_SG2M_, and *V*_birth_ are, respectively, cell cycle duration or the time between two divisions, time from division to next emergence of bud, time from onset of bud to next division, and volume of the cell at birth.

|  | Mother Cell : (mean , CV) |  | Daughter Cell : (mean , CV) |  |
| --- | --- | --- | --- | --- |
|  | Hybrid model | Experiment | Hybrid model | Experiment |
| $T_{\text{div}}$ (min) | (87.33 , 0.26) | (87 , 0.14) | (111.66 , 0.33) | (112.0 , 0.22) |
| $T_{\text{G1}}$ (min) | (18.23 , 0.36) | (16 , 0.5) | (37.51 , 0.59) | ( 37.0 , 0.5) |
| $T_{\text{SG2M}}$ (min) | (69.10 , 0.30) | (72 , 0.17) | (74.15 , 0.36) | (76.0 , 0.2) |
| $V_{\text{birth}}$ (fL) | (40.89 , 0.28) | (40 , 0.18) | ( 27.26 , 0.28) | (28.0 , 0.20) |

Next, we compare our simulations to the observed distributions of mRNA molecules in wild-type yeast cells. Figure 2 shows the histograms of 10 unregulated (*mCdh*1, *mTem*1, *mCdc*15, *mCdc*14, *mNet*1, *mCdc*55, *mEsp*1, *mSBF*, *mMBF*, *mMcm*1, *mAPC*) and 8 transcriptionally regulated (*mClb*5, *mCln*2, *mClb*2, *mSic*1, *mCdc*6, *mSwi*5, *mCdc*20, *mPds*1) mRNAs, in comparison to the distributions measured by Ball *et al.* [28]. Generally, there is a good agreement between the observed and simulated mRNA distributions, except for *mCln*1 + *mCln*2 and *mCdc*20. As expected, the unregulated transcripts follow Poisson distributions, which are consistent with experimental measurements. The cell-cycle regulated transcripts, which follow long-tailed, non-Poisson distributions, are well-fit by two-component Poisson distributions as reported by [26, 28]. (We note that in our model *mClb*2 represents both *mClb*1 and *mClb*2, and *mCln*2 = *mCln*1 + *mCln*2, whereas in the experiment these cyclin mRNAs are tracked independently. Therefore, we do not expect a particularly good match between the computed and observed distributions for these transcripts.)

**Figure 2:**
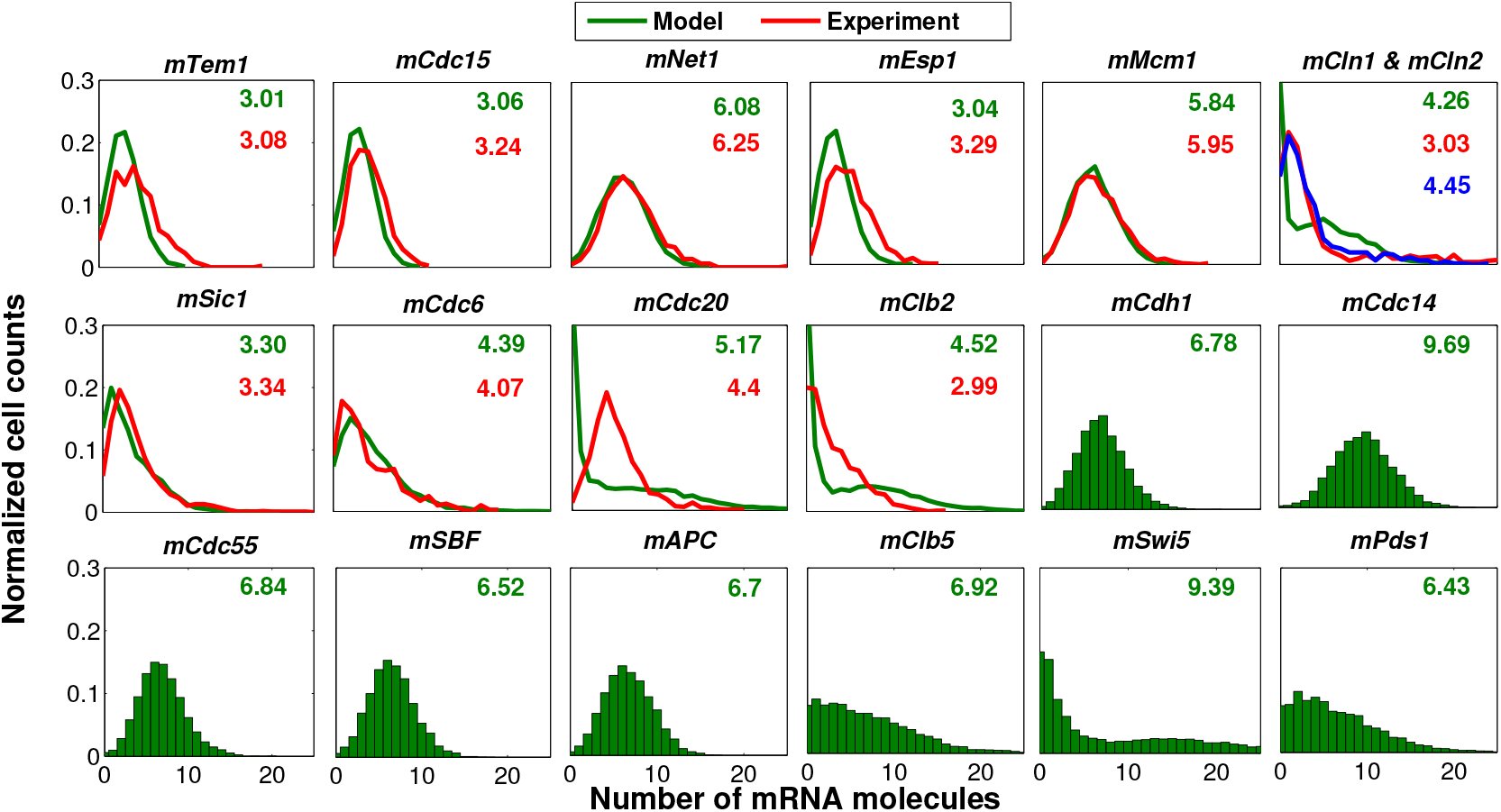
Histograms of mRNAs for a population of wild-type cells growing in glucose medium. The histograms of mRNA molecules generated from a stochastic run of the hybrid model (in green) are compared with experimental observations [28] (in red and blue colors) for a population of wild-type cells growing in glucose. (In the simulation the growth rate is set to 0.0072 min^−1^ to reproduce the 96 min mass-doubling time of wild-type cells growing in glucose culture medium.) The histograms in red are reproduced from the experimental data reported by Ball *et al.*[28]. The average number of mRNA molecules is reported on the histogram and compared with experiment where available. In our model *mCln*2 stands for both *mCln*1 + *mCln*2. In experiment, however, they are measured separately. Here, the histograms in red and blue are, respectively, *mCln*1 and *mCln*2. Similarly, in our model *mClb*2 describes the abundances of both *mClb*1 and *mClb*2. However, the histogram reproduced from the experimental data refers only to *mClb*2.

Table 2 compares the average abundances of proteins as observed in [51] and simulated by our model. The agreement is quite good, except for the Sic1.

**Table 2:** Average abundances of protein molecules per cell. The average abundance of protein molecules per cell, calculated from a simulation of the hybrid stochastic model, is compared with experimental observations reported in [51]. In our model Clb5 stands for Clb5 and Clb6, Clb2 stands for Clb1 and Clb2, and Cln2 stands for Cln1 and Cln2. We are reporting the total abundance of each protein, which includes protein molecules that are either phosphorylated or unphosphorylated, and that are bound in complexes or free. That is, Clb5_T_ = Clb5 + C5 + C5P + F5 + F5P, Clb2_T_ = Clb2 + C2 + C2P + F2 + F2P.

| Protein | Average abundance |  | Protein | Average abundance |  |
| --- | --- | --- | --- | --- | --- |
|  | Experiment | Hybrid model |  | Experiment | Hybrid model |
| **Cln3** | 108 | 128 | **Swi5<sub>T</sub>** | 688 | 661 |
| **Cln2<sub>T</sub>**<br>**= Cln1 + Cln2** | 1589<br>= 319 + 1270 | 1527 | **Tem1** | 573 | 542 |
| **Clb5<sub>T</sub>** | 420 | 463 | **Cdc15** | 238 | 245 |
| **Clb2<sub>T</sub>** | 693 | 700 | **Net1<sub>T</sub>** | 1590 | 1810 |
| **Sic1<sub>T</sub>** | 768 | 471 | **Cdc55** | 3170 | 3203 |

### 3.2 Mutant Cases

#### 3.2.1 Viability Criteria

Our hybrid stochastic model is used to simulate 122 mutant strains listed in Supplementary Table S5 in Supplementary Text. Prior to presenting simulation results of these mutants, we discuss the criteria for determining viability in a general stochastic model.

In the original deterministic model, a cell is considered viable if the following conditions are met:

1. certain events, listed in [36], take place in a proper sequence,
2. in particular, cell division occurs after budding, and
3. cell mass does not exceed a predetermined threshold (*mass* at division < 10).

The same set of rules should also apply to the stochastic model. A division is considered successful if the aforementioned viability criteria are met; otherwise, the cell is considered to be inviable. From our numerical simulations, the probability of successful division *p* is estimated by

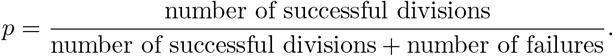

Using this metric, we consider a mutant strain viable if *p* > 0.75, inviable if *p* < 0.65, and partially viable otherwise. This viability criterion is based on the following considerations.

Let *N*_0_ be the initial population of cells in an experiment. After one cycle, the average number of cells that divide is *pN*_0_, while (1 − *p*)*N*_0_ cells exit the cycle and stop dividing. Thus, after one cycle, the total population of cells is (1 + *p*)*N*_0_, of which 2*pN*_0_ cells completed the previous cell cycle and (1 − *p*)*N*_0_ cells have ceased to divide (we call them dead cells). In our simulations, we disregard the (1 − *p*)*N*_0_ dead cells; hence, the number of actively dividing cells in the second cycle is *N*_1_ = 2*pN*_0_, and the expected number of actively dividing cells after *k* cycles will be *N_k_* = (2*p*)^*k*^N_0_.

In cell-viability experiments, colony formation is typically assessed after 24 h growth of a series of ten-fold diluted inocula. For wild-type yeast cells (*p* ≈ 1) growing on rich glucose medium (cycle time ≈ 12 h), each inoculum should increase by a factor of about 2^12^ = 4096. The colony sizes after 24 h growth of ten-fold serial dilutions would be (4000 *N*_0_, 400 *N*_0_, 40 *N*_0_, 4 *N*_0_), of which the first would be too dense to quantify, the last would be too sparse to see, and the middle two would be used to assess viability of mutant strains. For a mutant cell with *p* < 0.5, no visible colony will grow from the initial inoculum, and the mutant will be scored ‘inviable’. For mutant strains with 0.5 < *p* < 1, we must consider how the colony growth assay compares to wild-type cells. For *p* = 0.8 the initial inoculum grows to 280 *N*_0_, which is comparable to the first dilution of the wild-type cells, and we would score this mutant strain as ‘viable’. For *p* = 0.75 the initial inoculum grows to 130 *N*_0_, which is denser than the second dilution of the wild-type cells, and we would score this mutant strain as ‘probably viable’. For *p* = 0.65 the initial inoculum grows to 23 *N*_0_, which is less dense than the second dilution of the wild-type cells, and we would score this mutant strain as ‘hardly viable’. These calculations suggest that a mutant strain be considered viable if *p* ≥ 0.75 and nonviable if *p* ≤ 0.65. For 0.65 < *p* < 0.75, the strain is identified as partially viable.

Based on these criteria, we assessed the viability of 122 mutant strains of budding yeast that were studied in the modeling paper of Chen *et al.* [11]. To demonstrate the significant roles of noise in some of these mutants, we discuss two multiple-mutant strains, *cln*1Δ *cln*2Δ *bck*2Δ and *cln*3Δ *bck*2Δ multi-copy *CLN* 2 (Fig. 3), in some detail. According to experimental observations, the *cln*1Δ *cln*2Δ *bck*2Δ strain [60] is viable. However, due to deletion of START cyclins Cln1 and Cln2, the cell requires a longer time than normal to form a bud and hence grows to a larger size at division, in comparison with wild-type cells. Figure 3A shows that in the deterministic model the cell consistently exits mitosis and divides successfully with size larger than normal, as observed experimentally [60]. In the hybrid stochastic simulation, however, due to the stochastic nature of the process there is a finite probability that a cell may exit the cycle and become arrested in some phase of the cell cycle. In Fig. 3B for instance the cell grows too large in G1 phase and never divides again, while in Fig. 3C, it exits mitosis and divides successfully. The probability of successful completion of the cell division cycle, in this case, is computed to be *p* ≈ 0.84. As shown in Fig. 3D, the total number of cells in our computational culture increases exponentially, with a number-doubling time (NDT) of 140 min, which is slower than the number-doubling time NDT (approx 100 min) of a fully viable wild-type culture. Therefore, we conclude that the hybrid stochastic simulation correctly confirms the viability, but the reduced growth rate, of the *cln*1Δ *cln*2Δ *bck*2Δ strain.

**Figure 3:**
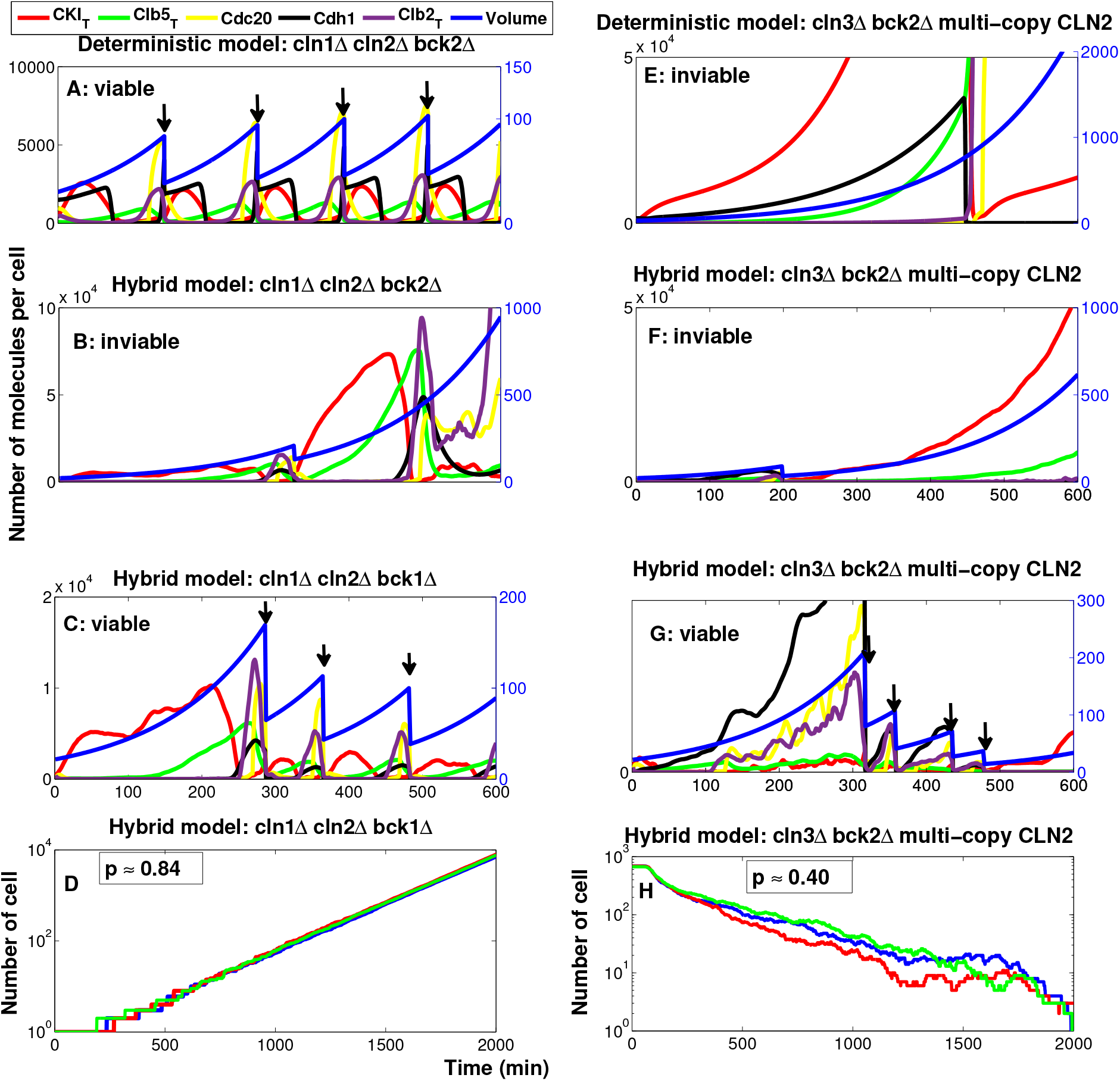
Comparison of deterministic and stochastic trajectories in two different double-mutant strains. **A:** Deterministic trajectories of *cln*1Δ *cln*2Δ *bck*2Δ; the cell consistently exits mitosis and divides (the divisions are indicated by arrows). **B-C:** Stochastic trajectories of *cln*1Δ *cln*2Δ *bck*2Δ from two independent runs. In panel B the cell becomes arrested in G1 phase while in panel C the cell divides successfully. **D:** The total number of cells as a function of time; we start each simulation with one cell and count the total number of cells over time for 2000 min. The probability of division is calculated as *p ≈* 0.84 which indicates that the *cln*1Δ *cln*2Δ *bck*2Δ strain is viable according to our definition. The semilog plot in panel D shows that the number of cells increases exponentially (NDT ≈ 140 min) in our computational culture. **E:** Deterministic trajectories of *cln*3Δ *bck*2Δ multi-copy *CLN* 2; the cell arrests permanently in G1 phase. **F-G:** Stochastic trajectories of *cln*3Δ *bck*2Δ multi-copy *CLN* 2 from two independent runs. In panel F the cell becomes arrested in G1 phase after one cycle, while in panel G the cell exits mitosis and divides successfully several times. **H:** The total number of cells as a function of time; we start the simulation with 1000 cells and count the total number of viable cells over time for 2000 min. The probability of division is calculated as *p* ≈ 0.40 which indicates that the *cln*3Δ *bck*2Δ multi-copy *CLN* 2 strain is inviable. The semilog plot in panel H shows that the total number of cells decreases exponentially in our computational culture.

Next we consider the inviable mutant strain *cln*3Δ *bck*2Δ multi-copy *CLN* 2 [61]. Figure 3E shows that in the deterministic model the mutant cell is arrested in G1 phase and grows without dividing until it dies. In the hybrid stochastic simulation, although many of the cells become arrested in G1 (see Fig. 3F), some cells manage to exit G1 phase, complete the cell cycle, commence a new cycle and divide a few times (see Fig. 3G). Nonetheless, according to Fig. 3H the total number of cells in our computational culture declines with time, because the probability of cell division is only *p* ≈ 0.40. Therefore, we conclude that the hybrid stochastic simulation correctly confirms the inviability of *cln*3Δ *bck*2Δ multi-copy *CLN* 2 strain.

Based on our hybrid stochastic simulations of all 122 mutant strains in Chen’s data base, we find that the model successfully reproduces the phenotypes of 103 of these strains. Our results for all mutant strains are reported in Supplementary Table S5 in Supplementary Text.

*CLB*2*db*Δ *clb*5Δ is a mutant with an interesting stochastic phenotype: it is inviable when grown on glucose medium but ‘partially viable’ when grown on raffinose (a sugar that supports a slower growth rate than glucose) [62]. Due to deletion of the destruction box of *CLB*2, Clb2 protein is in excess at telophase and the cell is unable to exit mitosis and divide, even in the absence of Clb5 protein (due to deletion of the *CLB*5 gene). When growing on raffinose, however, many of these mutant cells (approximately 60%-75%) are able to exit mitosis and commence a new cycle, whereas the remaining cells (25%-40%) are arrested in telophase and never re-enter the cell cycle [21]. In fact, the NDT of the double-mutant cells (250 - 300 min) is observed to be much longer than the NDT of wild-type cells (160 min) growing in raffinose medium [21].

Simulation results of Chen’s deterministic model predict that *CLB*2*db*Δ *clb*5Δ cells are inviable on glucose and viable on galactose and raffinose media. Clearly, we cannot expect a deterministic model to capture the stochastic properties of such a ‘partially viable’ mutant strain. Our hybrid stochastic model, however, describes the phenotype in exquisite detail. The probability of division for *CLB*2*db*Δ *clb*5Δ strain is *p* ≈ 0.65 confirming the partial viability of the mutant according to our viability criterion. Figure 4A shows the cumulative probability, *P* (*T*), of cycle times for wild-type and *CLB*2*db*Δ *clb*5Δ mutant cells growing in raffinose. (*P* (*T*) is the probability that the cycle time of a randomly chosen cell is longer than a specified time, *T*.) As shown in Fig. 4A, *P* (*T*) for the mutant cells levels off at approximately 35% as *T* increases, whereas, for wild-type cells, *P* (*T*) drops steadily (below 5%) as *T* increases beyond 250 min. Cumulative distributions of cycle times computed by our hybrid stochastic model (black lines) are in excellent agreement with the experimental distributions (red-blue-green lines) for both wild-type and the double-mutant cells.

**Figure 4:**
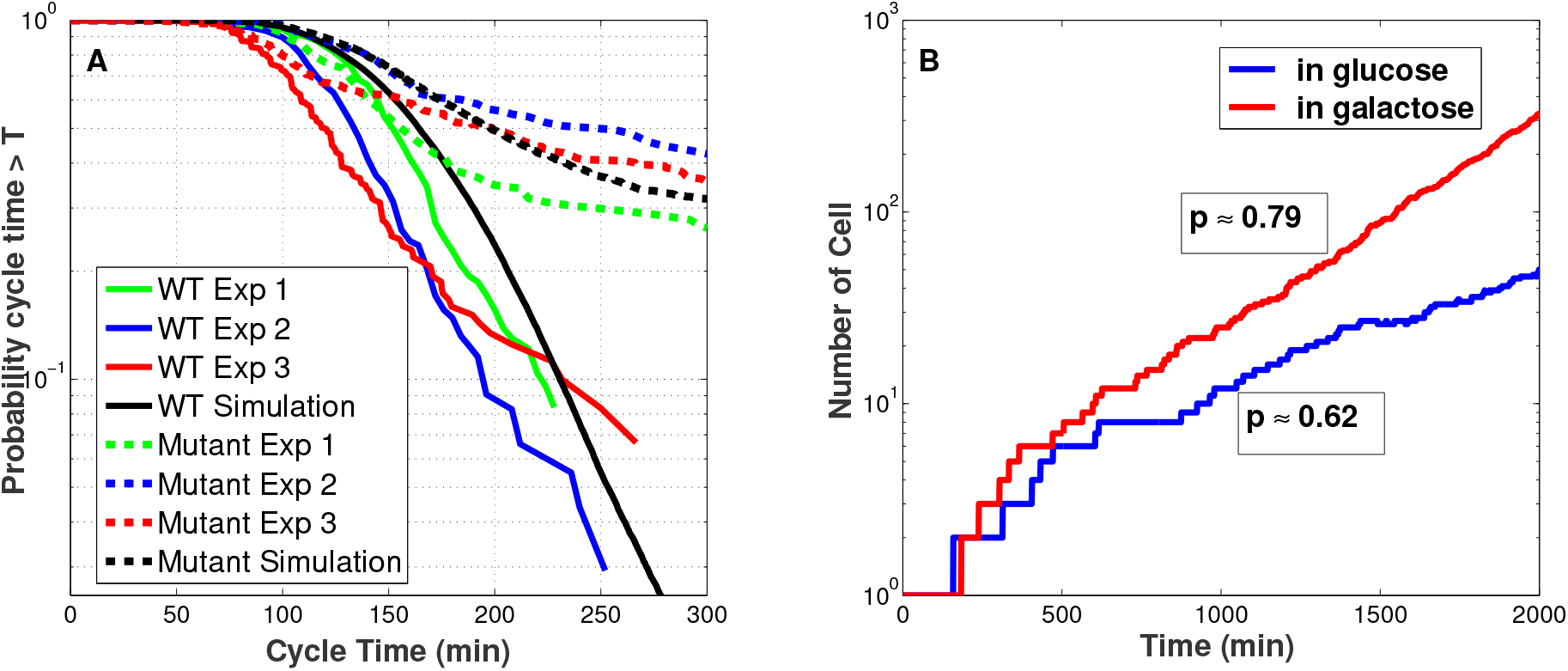
Stochastic phenotypes of two more mutant strains. **A:** Comparison of wild-type and *CLB*2*db*Δ *clb*5Δ mutant cells growing in raffinose. The probability that a cell divides with a cycle time longer than a specific time *T* is plotted for wild-type cells (solid lines) and mutant cells (dotted lines). The black lines are generated by our hybrid stochastic model and the red-blue-green lines are the results of three independent experimental runs by Ball *et al.* [21]. To model growth on raffinose medium in our simulation, the specific growth rate of cells is set to 0.00433 min^−1^ (MDT = 160 min). **B:** Comparison of cell proliferation for colonies of *CLB*1 *clb*2Δ *cdh*1Δ cells growing in glucose (blue) or galactose (red). The probability of division in our computational culture is given in the boxes next to each simulation. To mimic growth in glucose and galactose media, respectively, the specific growth rates are set to 0.0072 and 0.004621 min^−1^, i.e., MDT = 96 and 150 min, respectively.

Another interesting mutant strain is *CLB*1 *clb*2Δ *cdh*1Δ, for which the *CLB*1 gene is intact and *CLB*2 and *CDH*1 genes are deleted. Due to the mutual antagonism between Clb2 and Cdh1, deletion of both genes, *CDH*1 and *CLB*2, might be consistent with viability of the double-mutant strain, provided Clb1 is still functional. Indeed, experimental observations show that *CLB*1 *clb*2Δ *cdh*1Δ cells are *poorly viable* in glucose medium and *viable* when growing on galactose [62]. Chen’s deterministic model does not capture this phenotype; the model predicts the mutant cells to be viable in both media. However, simulation results of our hybrid stochastic model (Figure 4B) can reproduce the observed phenotype. The probability of division computed for a population of cells growing in glucose is ≈ 0.62, which suggests *poor* viability. In galactose the corresponding probability of division is ≈ 0.79, which indicates that the mutant grows well in this medium. Figure 4B confirms a faster increase in cell number in the slower growth medium (galactose) which is in agreement with experimental observations.

## 4 Discussion

In this paper we present a hybrid stochastic model of the molecular mechanism controlling progression through the budding yeast cell cycle. Our model provides a good match with experimental observations of many important characteristics of the budding yeast cell cycle, including inter-division time, cell size, and the phenotypes of more than 100 mutant strains. Compared with other approaches to stochastic modeling, our hybrid stochastic approach has several advantages. In a multi-scale regulatory network such as cell cycle controls, the major source of intrinsic noise can be attributed to low copy numbers of mRNA species in the gene-protein regulatory network. In fact, in budding yeast cells there are only 5-10 copies of each mRNA species encoding the production of corresponding proteins at levels of 500-5,000 molecules per cell. In such circumstances, small fluctuations in the population of mRNAs will result in substantial fluctuations in the corresponding protein levels. With this in mind, the key idea of the hybrid scheme is to partition the dynamics of mRNA species into the stochastic regime, in order to capture the major effects of random fluctuations in mRNA numbers, and to keep the protein dynamics in the deterministic framework, to achieve greater simulation efficiency. In addition, in this scheme it is not necessary to reformulate the complex rate laws governing protein interactions as elementary mass-action rate laws, which is a great advantage from a modeling standpoint.

In this paper, we have applied our hybrid stochastic method to a detailed molecular mechanism of cell cycle controls in budding yeast [11]. To apply our scheme to Chen’s model, which is a deterministic model of protein interactions, we first had to extend the model to include mRNA species that are transcribed from cell-cycle genes and translated into proteins. Then we carried out comprehensive simulations of wild-type yeast cells and more than 100 mutant strains, using both the deterministic and hybrid ODE/SSA models. Our stochastic model predicts the statistical properties of many different cell-cycle variables, including inter-division times, size at birth, and the abundances of specific mRNAs and proteins, and our stochastic simulations are in accord with most experimental observations, including detailed phenotypic characteristics of 103 out of 122 mutant cases. (Although Chen’s deterministic model may seem to ‘score’ better on the ‘viability’ of mutant strains, it is not so highly constrained as our stochastic model by consideration of the statistical properties of these mutant cells, especially the characteristics of ‘partially viable’ mutants.) In addition, our results prove that our hybrid approach to stochastic/deterministic simulations can achieve a good trade-off between accuracy and efficiency of numerical simulations. FORTRAN code takes about 15 min to simulate 10,000 cell cycle on an Intel i7-3770 processor with 16G memory running a Linux environment. A similar system using a fully stochastic model may take more than one day (the FORTRAN code by Barik *et al.*[26] is run using the same work station; it takes more than 30 hours to generate similar computational population of yeast cells).

## Supplementary Information

Supplementary information includes two files: one for the Supplementary Text and one for the Supplementary Code. In the Supplementary Text, we present more details for the cell cycle model used in this paper. Supplementary Tables S1-S4 list the time-dependent variables, differential equations, reactions and propensity functions, and parameter values. In Supplementary Table S5 we compare simulation results of 122 mutant strains with the observed phenotypes in experiment. The Supplementary Code file includes our hybrid model code in FORTRAN and statistical analysis code in MATLAB.

## Acknowledgments

This work was partially supported by the National Science Foundation under awards CCF-0953590, CCF-1526666, and MCB-1613741. Dr. Peccoud and Dr. Tyson’s work was also partially supported by NIH under award GM078989.

## Author contributions

M.A. and Y.C conceived of the presented idea. M.A. developed the simulation code and performed the computations. J.P provided the experimental data. Y.C. and J.J.T verified the analytical methods and supervised the findings of this work. All authors discussed the results and contributed to the final manuscript.

## Data availability

The datasets generated and analyzed during the current study are reproducible using the FORTRAN and MATLAB codes provided in the Supplementary Code. All other desired data sets are available from the corresponding author upon request.

