## Supplementary Text for "A Hybrid Stochastic Model of the Budding Yeast Cell Cycle"

### Cell cycle model

The main regulator of the cell cycle in budding yeast (Cdc28) is a cyclin-dependent protein kinase, which is constitutively expressed. Cdc28 forms an active kinase by binding to two families of cyclin partners, Cln1-3 and Clb1-6, each of which is synthesized and degraded in different phases of the cell cycle. In early G1 phase, Cln3 is the only available partner of Cdc28. As a newborn cell grows, the hetero-dimer of Cln3:Cdc28 accumulates until the cell reaches a critical size, when there is sufficient Cln3:Cdc28 (and a back-up protein, Bck2) to activate the transcription factors SBF and MBF. These transcription factors are responsible for the production of Cln2 and Clb5, respectively. Cln2 accumulation induces emergence of the bud and Clb5 initiates DNA synthesis.

In G1 phase the cyclin-dependent kinase inhibitors (CKI), Sic1 and Cdc6, are in high abundance, and they inhibit the activity of Clb5:Cdc28. However, as the activity of Cln2:Cdc28 increases, these inhibitors get phosphorylated and degraded by the SCF-proteosome pathway. As a result Clb5 accumulates to initiate DNA synthesis (S phase). Clb2 is the cyclin responsible for driving the cell into mitosis (M phase). As CKIs are removed, Clb2 level rises because Clb2 activates its own transcription factor Mcm1 in an autocatalytic reaction. Rising Clb2:Cdc28 activity phosphorylates and inactivates the transcription factors SBF and MBF. In telophase, Clb2 must be degraded below a threshold so that the cell can exit mitosis and return to G1 phase. Clb2 degradation is initiated by two proteins, Cdc20 and Cdh1. Cdc20 has been kept inactive in the early stages of mitosis by the 'mitotic checkpoint complex'. When all chromosomes are properly aligned on the mitotic spindle, Cdc20 becomes active and facilitates the degradation of Clb2.

Moreover, as a yeast cell exits mitosis, Cdh1 is activated by a phosphatase, Cdc14, which has been sequestered in the nucleolus by binding to Net1. After full chromosome alignment on the metaphase plate, first Tem1 and then Cdc15 become active. Cdc15 phosphorylates Net1, which leads to the release of Cdc14. Next, Cdc14 activates Cdh1 (which takes over for Cdc20 as the primary initiator of Clb1-6 degradation in G1), and Cdc14 also activates the transcription factor, Swi5, for production of CKIs in G1 phase. In this way the scene is set for the cell to return to G1 phase, when the CKIs are abundant and all cyclins (except for Cln3) are out of the picture.

### Variables, equations, reactions and parameter values

Supplementary Table S1. Variables and initial values. Except for mass which is a normalized (dimensionless) variable, all other variables are based on number of molecules per cell. Since initial values of variables expressed in algebraic equations are dependent of the variables expressed in ODEs, no initial values are listed for those variables.

| Variable | Description | Initial value |
|---|---|---|
| mass | Indicator of cell size (dimensionless) | 1.04 |
| Cln2 | Total cyclins Cln1 and Cln2 (represented as Cln2 in the model) | 21 |
| Clb5 | Active forms of cyclins Clb5 and Clb6 (represented as Clb5 in the model) | 48 |
| Clb2 | Active forms of Clb1 and Clb2 (represented as Clb2 in the model) | 233 |
| Sic1 | A stoichiometric inhibitor of Cdc28/Clb2 and Cdc28/Clb5 | 8 |
| Sic1P | Phophorylated form of Sic1 | 6 |
| C2 | A complex formed by Clb2 and Sic1 | 149 |
| C5 | A complex formed by Clb5 and Sic1 | 32 |
| C2P | Phophorylated form of C2 | 31 |
| C5P | Phophorylated form of C5 | 6 |
| Cdc6 | A stoichiometric inhibitor of Cdc28/Clb2 | 22 |
| Cdc6P | Phophorylated form of Cdc6 | 11 |
| F2 | A complex formed by Clb2 and Cdc6 | 66 |
| F5 | A complex formed by Clb5 and Cdc6 | 0 |
| F2P | Phophorylated form of F2 | 26 |
| F5P | Phophorylated form of F5 | 0 |

| | | |
|---|---|---|
| Swi5 | Transcription factor for Sic1 and Cdc6 | 613 |
| $Swi5_T$ | Total Swi5 | 634 |
| APC | Anaphase Promoting Complex | 285 |
| APCP | Active (phosphorylated) form of APC | 16 |
| $Cdc20_T$ | Total Cdc20 | 4059 |
| Cdc20A | Active form of Cdc20 (an APC partner) | 752 |
| Cdh1T | Total Cdh1 | 1273 |
| Cdh1 | Active Cdh1 (an APC partner) | 895 |
| Tem1 | A GTP-binding protein | 1140 |
| $Tem1_T$ | Total Tem1 | 1166 |
| Cdc15 | A kinase required for nuclear division | 342 |
| $Cdc15_T$ | Total Cdc15 | 518 |
| Cdc14 | A phosphatase required for mitosis exit | 121 |
| $Cdc14_T$ | Total of Cdc14 | 607 |
| Net1 | A stoichiometric inhibitor of Cdc14 | 7 |
| $Net1_T$ | Total Net1 | 838 |
| Net1P | Phosphorylated form of Net1 | - |
| RENT | A protein complex of Net1 and Cdc14 that regulates nucleolar silencing and telophase | 324 |
| RENTP | Phosphorylated form of RENT | - |
| Cdc55 | Active form of CDC55 phosphatase | 1161 |
| Esp1 | A protein required for sister chromatid separation | 9 |
| $Esp1_T$ | Total Esp1 | 42 |
| Pds1 | A stoichiometric inhibitor of Esp1 | 2 |

| | | |
|---|---|---|
| ORI | An auxiliary variable representing proteins that signal the onset of DNA synthesis | 2374 |
| BUD | An auxiliary variable representing proteins that initiate a new bud | 0 |
| SPN | An auxiliary variable representing proteins that signals the alignment of all chromosomes | 0 |
| SBF | Transcription factor for Cln2 | - |
| $SBF_T$ | Total SBF | 527 |
| MBF | Transcription factor for Clb5 | - |
| $MBF_T$ | Total MBF | 442 |
| Mcm1 | Transcription factor for Clb2, Cdc20 and Swi5 | - |
| $Mcm1_T$ | Total Mcm1 | 10363 |
| Cln3 | A G1-cyclin initiating START events | 50 |
| Bck2 | A back-up protein initiating START events | 25 |
| $Clb5_T$ | Total Clb5 | - |
| $Clb2_T$ | Total Clb2 | - |
| $Sic1_T$ | Total Sic1 | - |
| $Cdc6_T$ | Total Cdc6 | - |
| $CKI_T$ | Total cyclin inhibitors Sic1 and Cdc6 | - |
| $mCdh1$ | Cdh1 messenger RNA | 4 |
| $mTem1$ | Tem1 messenger RNA | 2 |
| $mCdc15$ | Cdc15 messenger RNA | 2 |
| $mCdc14$ | Cdc14 messenger RNA | 6 |
| $mNet1$ | Net1 messenger RNA | 3 |

| | | |
|---|---|---|
| $mCdc55$ | Cdc55 messenger RNA | 4 |
| $mEsp1$ | Esp1 messenger RNA | 2 |
| $mSBF$ | SBF messenger RNA | 4 |
| $mMBF$ | MBF messenger RNA | 4 |
| $mMcm1$ | Mcm1 messenger RNA | 3 |
| $mAPC$ | APC messenger RNA | 4 |
| $mCln2$ | Cln2 messenger RNA | 1 |
| $mClb5$ | Clb5 messenger RNA | 1 |
| $mClb2$ | Clb2 messenger RNA | 4 |
| $mSic1$ | Sic1 messenger RNA | 3 |
| $mCdc6$ | Cdc6 messenger RNA | 4 |
| $mSwi5$ | Swi5 messenger RNA | 9 |
| $mCdc20$ | Cdc20 messenger RNA | 5 |
| $mPds1$ | Pds1 messenger RNA | 4 |

Supplementary Table S2. Equations.

| |
|---|
| $\frac{\mathrm{d}mass}{\mathrm{d}t} = k_{\mathrm{g}} \cdot mass$ |
| $\frac{\mathrm{d}Cln2}{\mathrm{d}t} = (k'_{\mathrm{s,n2}} \cdot c_{\mathrm{cln2}} \cdot mass + k''_{\mathrm{s,n2}} \cdot \frac{c_{\mathrm{cln2}}}{c_{\mathrm{sbf}}} \cdot SBF) \cdot mass \cdot mCln2 - k_{\mathrm{d,n2}} \cdot Cln2 + k_{\mathrm{g}} \cdot Cln2$ |
| $\frac{\mathrm{d}Clb5}{\mathrm{d}t} = (k'_{\mathrm{s,b5}} \cdot c_{\mathrm{clb5}} \cdot mass + k''_{\mathrm{s,b5}} \cdot \frac{c_{\mathrm{clb5}}}{c_{\mathrm{mbf}}} \cdot MBF) \cdot mass \cdot mClb5 + (k_{\mathrm{d3,c1}} \cdot C5\mathrm{P} + k_{\mathrm{di,b5}} \cdot C5) + (k_{\mathrm{d3,f6}} \cdot F5\mathrm{P} + k_{\mathrm{di,f5}} \cdot F5) - (V_{\mathrm{d,b5}} + \frac{k_{\mathrm{as,b5}}}{c_{\mathrm{sic1}} \cdot mass} \cdot Sic1 + \frac{k_{\mathrm{as,f5}}}{c_{\mathrm{cdc6}} \cdot mass} \cdot Cdc6) \cdot Clb5 + k_{\mathrm{g}} \cdot Clb5$ |
| $\frac{\mathrm{d}Clb2}{\mathrm{d}t} = (k'_{\mathrm{s,b2}} \cdot c_{\mathrm{clb2}} \cdot mass + k''_{\mathrm{s,b2}} \cdot \frac{c_{\mathrm{clb2}}}{c_{\mathrm{mcm1}}} \cdot Mcm1) \cdot mass \cdot mClb2 + (k_{\mathrm{d3,c1}} \cdot C2\mathrm{P} + k_{\mathrm{di,b2}} \cdot C2) + (k_{\mathrm{d3,f6}} \cdot F2\mathrm{P} + k_{\mathrm{di,f2}} \cdot F2) - (V_{\mathrm{d,b2}} + \frac{k_{\mathrm{as,b2}}}{c_{\mathrm{sic1}} \cdot mass} \cdot Sic1 + \frac{k_{\mathrm{as,f2}}}{c_{\mathrm{cdc6}} \cdot mass} \cdot Cdc6) \cdot Clb2 + k_{\mathrm{g}} \cdot Clb2$ |
| $\frac{\mathrm{d}Sic1}{\mathrm{d}t} = (k'_{\mathrm{s,c1}} \cdot c_{\mathrm{sic1}} \cdot mass + k''_{\mathrm{s,c1}} \frac{c_{\mathrm{sic1}}}{c_{\mathrm{swi5}}} \cdot Swi5) \cdot mSic1 + (V_{\mathrm{d,b2}} + k_{\mathrm{di,b2}}) \cdot C2\mathrm{P} + (V_{\mathrm{d,b5}} + k_{\mathrm{di,b5}}) \cdot C5\mathrm{P} + \frac{k_{\mathrm{pp,c1}}}{c_{\mathrm{cdc14}} \cdot mass} \cdot Cdc14 \cdot Sic1\mathrm{P} - (\frac{k_{\mathrm{as,b2}}}{c_{\mathrm{clb2}} \cdot mass} \cdot Clb2 + \frac{k_{\mathrm{as,b5}}}{c_{\mathrm{clb5}} \cdot mass} \cdot Clb5 + V_{\mathrm{kp,c1}}) \cdot Sic1 + k_{\mathrm{g}} \cdot Sic1$ |
| $\frac{\mathrm{d}Sic1P}{\mathrm{d}t} = V_{\mathrm{kp,c1}} \cdot Sic1 - (\frac{k_{\mathrm{pp,c1}}}{c_{\mathrm{cdc14}} \cdot mass} \cdot Cdc14 + k_{\mathrm{d3,c1}}) \cdot Sic1\mathrm{P} + V_{\mathrm{d,b2}} \cdot C2\mathrm{P} + V_{\mathrm{d,b5}} \cdot C5\mathrm{P} + k_{\mathrm{g}} \cdot Sic1\mathrm{P}$ |
| $\frac{\mathrm{d}C2}{\mathrm{d}t} = \frac{k_{\mathrm{as,b2}}}{c_{\mathrm{clb2}} \cdot mass} \cdot \mathrm{Clb2} \cdot Sic1 + \frac{k_{\mathrm{pp,c1}}}{c_{\mathrm{cdc14}} \cdot mass} \cdot Cdc14 \cdot C2\mathrm{P} - (k_{\mathrm{di,b2}} + V_{\mathrm{d,b2}} + V_{\mathrm{kp,c1}}) \cdot C2 + k_{\mathrm{g}} \cdot C2$ |
| $\frac{\mathrm{d}C5}{\mathrm{d}t} = \frac{k_{\mathrm{as,b5}}}{c_{\mathrm{clb5}} \cdot mass} \cdot \mathrm{Clb5} \cdot Sic1 + \frac{k_{\mathrm{pp,c1}}}{c_{\mathrm{cdc14}} \cdot mass} \cdot Cdc14 \cdot C5\mathrm{P} - (k_{\mathrm{di,b5}} + V_{\mathrm{d,b5}} + V_{\mathrm{kp,c1}}) \cdot C5 + k_{\mathrm{g}} \cdot C5$ |
| $\frac{\mathrm{d}C2\mathrm{P}}{\mathrm{d}t} = V_{\mathrm{kp,c1}} \cdot C2 - (\frac{k_{\mathrm{pp,c1}}}{c_{\mathrm{cdc14}} \cdot mass} \cdot Cdc14 + k_{\mathrm{d3,c1}} + V_{\mathrm{d,b2}}) \cdot C2\mathrm{P} + k_{\mathrm{g}} \cdot C2\mathrm{P}$ |
| $\frac{\mathrm{d}C5\mathrm{P}}{\mathrm{d}t} = V_{\mathrm{kp,c1}} \cdot C5 - (\frac{k_{\mathrm{pp,c1}}}{c_{\mathrm{cdc14}} \cdot mass} \cdot Cdc14 + k_{\mathrm{d3,c1}} + V_{\mathrm{d,b5}}) \cdot C5\mathrm{P} + k_{\mathrm{g}} \cdot C5\mathrm{P}$ |
| $\frac{\mathrm{d}Cdc6}{\mathrm{d}t} = (k'_{\mathrm{s,f6}} \cdot c_{\mathrm{cdc6}} \cdot mass + k''_{\mathrm{s,f6}} \cdot \frac{c_{\mathrm{cdc6}}}{c_{\mathrm{swi5}}} \cdot Swi5 + k'''_{\mathrm{s,f6}} \cdot \frac{c_{\mathrm{cdc6}}}{c_{\mathrm{sbf}}} \cdot SBF) \cdot mCdc6 + (V_{\mathrm{d,b2}} + k_{\mathrm{di,f2}}) \cdot F2 + (V_{\mathrm{d,b5}} + k_{\mathrm{di,f5}}) \cdot F5 + \frac{k_{\mathrm{pp,f6}}}{c_{\mathrm{cdc14}} \cdot mass} \cdot Cdc14 \cdot Cdc6P - (\frac{k_{\mathrm{as,f2}}}{c_{\mathrm{clb2}} \cdot mass} \cdot Clb2 + \frac{k_{\mathrm{as,f5}}}{c_{\mathrm{clb5}} \cdot mass} \cdot Clb5 + V_{\mathrm{kp,f6}}) \cdot Cdc6 + k_{\mathrm{g}} \cdot Cdc6$ |
| $\frac{\mathrm{d}Cdc6\mathrm{P}}{\mathrm{d}t} = V_{\mathrm{kp,f6}} \cdot Cdc6 - (\frac{k_{\mathrm{pp,f6}}}{c_{\mathrm{cdc14}} \cdot mass} \cdot Cdc14 + k_{\mathrm{d3,f6}}) \cdot Cdc6\mathrm{P} + V_{\mathrm{d,b2}} \cdot F2\mathrm{P} + V_{\mathrm{d,b5}} \cdot F5\mathrm{P} + k_{\mathrm{g}} \cdot Cdc6\mathrm{P}$ |
| $\frac{\mathrm{d}F2}{\mathrm{d}t} = \frac{k_{\mathrm{as,f2}}}{c_{\mathrm{clb2}} \cdot mass} \cdot \mathrm{Clb2} \cdot Cdc6 + \frac{k_{\mathrm{pp,f6}}}{c_{\mathrm{cdc14}} \cdot mass} \cdot Cdc14 \cdot F2\mathrm{P} - (k_{\mathrm{di,f2}} + V_{\mathrm{d,b2}} + V_{\mathrm{kp,f6}}) \cdot F2 + k_{\mathrm{g}} \cdot F2$ |
| $\frac{\mathrm{d}F5}{\mathrm{d}t} = \frac{k_{\mathrm{as,f5}}}{c_{\mathrm{clb5}} \cdot mass} \cdot Clb5 \cdot Cdc6 + \frac{k_{\mathrm{pp,f6}}}{c_{\mathrm{cdc14}} \cdot mass} \cdot Cdc14 \cdot F5\mathrm{P} - (k_{\mathrm{di,f5}} + V_{\mathrm{d,b5}} + V_{\mathrm{kp,f6}}) \cdot F5 + k_{\mathrm{g}} \cdot F5$ |
| $\frac{\mathrm{d}F2\mathrm{P}}{\mathrm{d}t} = V_{\mathrm{kp,f6}} \cdot F2 - (\frac{k_{\mathrm{pp,f6}}}{c_{\mathrm{cdc14}} \cdot mass} \cdot Cdc14 + k_{\mathrm{d3,f6}} + V_{\mathrm{d,b2}}) \cdot F2\mathrm{P} + k_{\mathrm{g}} \cdot F2\mathrm{P}$ |
| $\frac{\mathrm{d}F5\mathrm{P}}{\mathrm{d}t} = V_{\mathrm{kp,f6}} \cdot F5 - (\frac{k_{\mathrm{pp,f6}}}{c_{\mathrm{cdc14}} \cdot mass} \cdot Cdc14 + k_{\mathrm{d3,f6}} + V_{\mathrm{d,b5}}) \cdot F5\mathrm{P} + k_{\mathrm{g}} \cdot F5\mathrm{P}$ |
| $\frac{\mathrm{d}Swi5_{\mathrm{T}}}{\mathrm{d}t} = (k'_{\mathrm{s,swi}} \cdot c_{\mathrm{swi5}} \cdot mass + k''_{\mathrm{s,swi}} \cdot \frac{c_{\mathrm{swi5}}}{c_{\mathrm{mcm1}}} \cdot Mcm1) \cdot mSwi5 - k_{\mathrm{d,swi}} \cdot Swi5T + k_{\mathrm{g}} \cdot Swi5_{\mathrm{T}}$ |
| $\frac{\mathrm{d}Swi5}{\mathrm{d}t} = (k'_{\mathrm{s,swi}} \cdot c_{\mathrm{swi5}} \cdot mass + k''_{\mathrm{s,swi}} \cdot \frac{c_{\mathrm{swi5}}}{c_{\mathrm{mcm1}}} \cdot Mcm1) \cdot mSwi5 + \frac{k_{\mathrm{a,swi}}}{c_{\mathrm{cdc14}} \cdot mass} \cdot Cdc14 \cdot (Swi5_{\mathrm{T}} - Swi5) - (k_{\mathrm{d,swi}} + \frac{k_{\mathrm{i,swi}}}{c_{\mathrm{clb2}} \cdot mass} \cdot Clb2) \cdot Swi5 + k_{\mathrm{g}} \cdot Swi5$ |
| *(continued)* |

Supplementary Table S2. (continued)

| |
|---|
| $\frac{\mathrm{d}APC\mathrm{P}}{\mathrm{d}t} = \frac{k_{\mathrm{a,apc}} \cdot \frac{c_{\mathrm{apc}}}{c_{\mathrm{clb2}}} \cdot Clb2 \cdot (APC - APC\mathrm{P})}{J_{\mathrm{a,apc}} \cdot c_{\mathrm{apc}} \cdot mass + APC - APC\mathrm{P}} - \frac{k_{\mathrm{i,apc}} \cdot c_{\mathrm{apc}} \cdot mass \cdot APC\mathrm{P}}{J_{\mathrm{i,apc}} \cdot c_{\mathrm{apc}} \cdot mass + APC\mathrm{P}} + k_{\mathrm{g}} \cdot APC\mathrm{P}$ |
| $\frac{\mathrm{d}Cdc20_{\mathrm{T}}}{\mathrm{d}t} = (k'_{\mathrm{s,20}} \cdot c_{\mathrm{cdc20}} \cdot mass + k''_{\mathrm{s,20}} \cdot \frac{c_{\mathrm{cdc20}}}{c_{\mathrm{mcm1}}} \cdot Mcm1) \cdot mCdc20 - k_{\mathrm{d,20}} \cdot Cdc20_{\mathrm{T}} + k_{\mathrm{g}} \cdot Cdc20_{\mathrm{T}}$ |
| $\frac{\mathrm{d}Cdc20\mathrm{A}}{\mathrm{d}t} = (k'_{\mathrm{a,20}} + \frac{k''_{\mathrm{a,20}}}{c_{\mathrm{cdc20}} \cdot mass} \cdot APC\mathrm{P}) \cdot (Cdc20_{\mathrm{T}} - Cdc20\mathrm{A}) - (k_{\mathrm{mad2}} + k_{\mathrm{d,20}}) \cdot Cdc20\mathrm{A} + k_{\mathrm{g}} \cdot Cdc20\mathrm{A}$ |
| $\frac{\mathrm{d}Cdh1_{\mathrm{T}}}{\mathrm{d}t} = k_{\mathrm{s,cdh}} \cdot c_{\mathrm{cdh1}} \cdot mass \cdot mCdh1 - k_{\mathrm{d,cdh}} \cdot Cdh1_{\mathrm{T}} + k_{\mathrm{g}} \cdot Cdh1_{\mathrm{T}}$ |
| $\frac{\mathrm{d}Cdh1}{\mathrm{d}t} = k_{\mathrm{s,cdh}} \cdot c_{\mathrm{cdh1}} \cdot mass \cdot mCdh1 - k_{\mathrm{d,cdh}} \cdot Cdh1 + \frac{V_{\mathrm{a,cdh}} \cdot c_{\mathrm{cdh1}} \cdot mass \cdot (Cdh1_{\mathrm{T}} - Cdh1)}{J_{\mathrm{a,cdh}} \cdot c_{\mathrm{cdh1}} \cdot mass + Cdh1_{\mathrm{T}} - Cdh1} - \frac{V_{\mathrm{i,cdh}} \cdot c_{\mathrm{cdh1}} \cdot mass \cdot Cdh1}{J_{\mathrm{i,cdh}} \cdot c_{\mathrm{cdh1}} \cdot mass + Cdh1} + k_{\mathrm{g}} \cdot Cdh1$ |
| $\frac{\mathrm{d}Tem1}{\mathrm{d}t} = \frac{k_{\mathrm{lte1}} \cdot c_{\mathrm{tem1}} \cdot mass \cdot (Tem1_{\mathrm{T}} - Tem1)}{J_{\mathrm{a,tem}} \cdot c_{\mathrm{tem1}} \cdot mass + Tem1_{\mathrm{T}} - Tem1} - \frac{k_{\mathrm{bub2}} \cdot c_{\mathrm{tem1}} \cdot mass \cdot Tem1}{J_{\mathrm{i,tem}} \cdot c_{\mathrm{tem1}} \cdot mass + Tem1} + k_{\mathrm{g}} \cdot Tem1$ |
| $\frac{\mathrm{d}Cdc15}{\mathrm{d}t} = (\frac{k'_{\mathrm{a,15}}}{c_{\mathrm{tem1}} \cdot mass} \cdot (Tem1_{\mathrm{T}} - Tem1) + \frac{k''_{\mathrm{a,15}}}{c_{\mathrm{tem1}} \cdot mass} \cdot \mathrm{Tem1} + \frac{k'''_{\mathrm{a,15}}}{c_{\mathrm{cdc14}} \cdot mass} \cdot Cdc14) \cdot (Cdc15T - Cdc15) - k_{\mathrm{i,15}} \cdot Cdc15 + k_{\mathrm{g}} \cdot Cdc15$ |
| $\frac{\mathrm{d}Cdc14_{\mathrm{T}}}{\mathrm{d}t} = k_{\mathrm{s,14}} \cdot c_{\mathrm{cdc14}} \cdot mass \cdot mCdc14 - k_{\mathrm{d,14}} \cdot Cdc14_{\mathrm{T}} + k_{\mathrm{g}} \cdot Cdc14_{\mathrm{T}}$ |
| $\frac{\mathrm{d}Cdc14}{\mathrm{d}t} = k_{\mathrm{s,14}} \cdot c_{\mathrm{cdc14}} \cdot mass \cdot mCdc14 - k_{\mathrm{d,14}} \cdot Cdc14 + k_{\mathrm{d,net}} \cdot (RENT + RENT\mathrm{P}) + k_{\mathrm{di,rent}} \cdot RENT + k_{\mathrm{di,rentp}} \cdot RENT\mathrm{P} + k_{\mathrm{g}} \cdot Cdc14 - (\frac{k_{\mathrm{as,rent}}}{c_{\mathrm{net1}} \cdot mass} \cdot Net1 + \frac{k_{\mathrm{as,rentp}}}{c_{\mathrm{net1}} \cdot mass} \cdot Net1\mathrm{P}) \cdot Cdc14$ |
| $\frac{\mathrm{d}Net1_{\mathrm{T}}}{\mathrm{d}t} = k_{\mathrm{s,net}} \cdot c_{\mathrm{net1}} \cdot mass \cdot mNet1 - k_{\mathrm{d,net}} \cdot Net1_{\mathrm{T}} + k_{\mathrm{g}} \cdot Net1_{\mathrm{T}}$ |
| $\frac{\mathrm{d}Net1}{\mathrm{d}t} = k_{\mathrm{s,net}} \cdot c_{\mathrm{cdc14}} \cdot mass \cdot mNet1 - k_{\mathrm{d,net}} \cdot Net1 + k_{\mathrm{d,14}} \cdot RENT + k_{\mathrm{di,rent}} \cdot RENT - \frac{k_{\mathrm{as,rent}}}{c_{\mathrm{cdc14}} \cdot mass} \cdot Cdc14 \cdot Net1 + V_{\mathrm{pp,net}} \cdot Net1\mathrm{P} - V_{\mathrm{kp,net}} \cdot Net1 + k_{\mathrm{g}} \cdot Net1$ |
| $\frac{\mathrm{d}RENT}{\mathrm{d}t} = -(k_{\mathrm{d,14}} + k_{\mathrm{d,net}}) \cdot RENT - k_{\mathrm{di,rent}} \cdot RENT + \frac{k_{\mathrm{as,rent}}}{c_{\mathrm{cdc14}} \cdot mass} \cdot Cdc14 \cdot Net1 + V_{\mathrm{pp,net}} \cdot RENT\mathrm{P} - V_{\mathrm{kp,net}} \cdot RENT + k_{\mathrm{g}} \cdot RENT$ |
| $\frac{\mathrm{d}Cdc55}{\mathrm{d}t} = k_{\mathrm{s,55}} \cdot c_{\mathrm{cdc55}} \cdot mass \cdot mCdc55 - V_{\mathrm{d,55}} \cdot Cdc55 + k_{\mathrm{g}} \cdot Cdc55$ |
| $\frac{\mathrm{d}Pds1}{\mathrm{d}t} = (k'_{\mathrm{s,pds}} \cdot c_{\mathrm{pds1}} \cdot mass + k''_{\mathrm{s1,pds}} \cdot \frac{c_{\mathrm{pds1}}}{c_{\mathrm{sbf}}} \cdot SBF + k''_{\mathrm{s2,pds}} \cdot \frac{c_{\mathrm{pds1}}}{c_{\mathrm{mcm1}}} \cdot Mcm1) \cdot mPds1 + k_{\mathrm{di,esp}} \cdot PE - (V_{\mathrm{d,pds}} + \frac{k_{\mathrm{as,esp}}}{c_{\mathrm{esp1}} \cdot mass} \cdot Esp1) \cdot Pds1 + k_{\mathrm{g}} \cdot Pds1$ |
| $\frac{\mathrm{d}Esp1}{\mathrm{d}t} = -\frac{k_{\mathrm{as,esp}}}{c_{\mathrm{pds1}} \cdot mass} \cdot Pds1 \cdot Esp1 + (k_{\mathrm{di,esp}} + V_{\mathrm{d,pds}}) \cdot PE + k_{\mathrm{g}} \cdot Esp1$ |
| $\frac{\mathrm{d}ORI}{\mathrm{d}t} = k_{\mathrm{s,ori}} \cdot (e_{\mathrm{ori,b5}} \cdot \frac{c_{\mathrm{ori}}}{c_{\mathrm{clb5}}} \cdot Clb5 + e_{\mathrm{ori,b2}} \cdot \frac{c_{\mathrm{ori}}}{c_{\mathrm{clb2}}} \cdot Clb2) - k_{\mathrm{d,ori}} \cdot ORI + k_{\mathrm{g}} \cdot ORI$ |

*(continued)*

| Supplementary Table S2. (continued) |
| --- |
| $\frac{\mathrm{d}BUD}{\mathrm{d}t} = k_{\mathrm{s,bud}} \cdot (e_{\mathrm{bud,n2}} \cdot \frac{c_{\mathrm{bud}}}{c_{\mathrm{cln2}}} \cdot Cln2 + e_{\mathrm{bud,n3}} \cdot \frac{c_{\mathrm{bud}}}{c_{\mathrm{cln3}}} \cdot Cln3 + e_{\mathrm{bud,b5}} \cdot \frac{c_{\mathrm{bud}}}{c_{\mathrm{clb5}}} \cdot Clb5) - k_{\mathrm{d,bud}} \cdot BUD + k_{\mathrm{g}} \cdot BUD$ |
| $\frac{\mathrm{d}SPN}{\mathrm{d}t} = \frac{k_{\mathrm{s,spn}} \cdot c_{\mathrm{spn}} \cdot mass \cdot Clb2}{J_{\mathrm{spn}} \cdot c_{\mathrm{clb2}} \cdot mass + Clb2} - k_{\mathrm{d,spn}} \cdot SPN + k_{\mathrm{g}} \cdot SPN$ |
| $\frac{\mathrm{d}Tem1_{\mathrm{T}}}{\mathrm{d}t} = k_{\mathrm{s,tem1t}} \cdot c_{\mathrm{tem1}} \cdot mass \cdot mTem1 - k_{\mathrm{d,tem1t}} \cdot Tem1_{\mathrm{T}} + k_{\mathrm{g}} \cdot Tem1_{\mathrm{T}}$ |
| $\frac{\mathrm{d}Cdc15_{\mathrm{T}}}{\mathrm{d}t} = k_{\mathrm{s,cdc15t}} \cdot c_{\mathrm{cdc15}} \cdot mass \cdot mCdc15 - k_{\mathrm{d,cdc15t}} \cdot Cdc15_{\mathrm{T}} + k_{\mathrm{g}} \cdot Cdc15_{\mathrm{T}}$ |
| $\frac{\mathrm{d}Esp1_{\mathrm{T}}}{\mathrm{d}t} = k_{\mathrm{s,esp1t}} \cdot c_{\mathrm{esp1}} \cdot mass \cdot mEsp1 - k_{\mathrm{d,esp1t}} \cdot Esp1_{\mathrm{T}} + k_{\mathrm{g}} \cdot Esp1_{\mathrm{T}}$ |
| $\frac{\mathrm{d}SBF_{\mathrm{T}}}{\mathrm{d}t} = k_{\mathrm{s,sbft}} \cdot c_{\mathrm{sbf}} \cdot mass \cdot mSBF - k_{\mathrm{d,sbft}} \cdot SBF_{\mathrm{T}} + k_{\mathrm{g}} \cdot SBF_{\mathrm{T}}$ |
| $\frac{\mathrm{d}MBF_{\mathrm{T}}}{\mathrm{d}t} = k_{\mathrm{s,mbft}} \cdot c_{\mathrm{mbf}} \cdot mass \cdot mMBF - k_{\mathrm{d,mbft}} \cdot MBF_{\mathrm{T}} + k_{\mathrm{g}} \cdot MBF_{\mathrm{T}}$ |
| $\frac{\mathrm{d}Mcm1_{\mathrm{T}}}{\mathrm{d}t} = k_{\mathrm{s,mcm1t}} \cdot c_{\mathrm{mcm1}} \cdot mass \cdot mMcm1 - k_{\mathrm{d,mcm1t}} \cdot Mcm1_{\mathrm{T}} + k_{\mathrm{g}} \cdot Mcm1_{\mathrm{T}}$ |
| $\frac{\mathrm{d}APC}{\mathrm{d}t} = k_{\mathrm{s,apct}} \cdot c_{\mathrm{apc}} \cdot mass \cdot mAPC - k_{\mathrm{d,apct}} \cdot APC + k_{\mathrm{g}} \cdot APC$ |
| $\frac{\mathrm{d}Cln3}{\mathrm{d}t} = k_{\mathrm{s,n3}} \cdot c_{\mathrm{cln3}} \cdot mass^2 - k_{\mathrm{d,n3}} \cdot Cln3 + k_{\mathrm{g}} \cdot Cln3$ |
| $\frac{\mathrm{d}Bck2}{\mathrm{d}t} = k_{\mathrm{s,k2}} \cdot c_{\mathrm{bck2}} \cdot mass^2 - k_{\mathrm{d,k2}} \cdot Bck2 + k_{\mathrm{g}} \cdot Bck2$ |
| $G(V_{\mathrm{a}}, V_{\mathrm{i}}, J_{\mathrm{a}}, J_{\mathrm{i}}) = \frac{2 J_{\mathrm{i}} V_{\mathrm{a}}}{V_{\mathrm{i}} - V_{\mathrm{a}} + J_{\mathrm{a}} V_{\mathrm{i}} + J_{\mathrm{i}} V_{\mathrm{a}} + \sqrt{(V_{\mathrm{i}} - V_{\mathrm{a}} + J_{\mathrm{a}} V_{\mathrm{i}} + J_{\mathrm{i}} V_{\mathrm{a}})^2 - 4(V_{\mathrm{i}} - V_{\mathrm{a}}) J_{\mathrm{i}} V_{\mathrm{a}}}}$ |
| $SBF = G(V_{\mathrm{a,sbf}}, V_{\mathrm{i,sbf}}, J_{\mathrm{a,sbf}}, J_{\mathrm{i,sbf}}) \cdot SBF_{\mathrm{T}}$ |
| $MBF = G(V_{\mathrm{a,sbf}}, V_{\mathrm{i,sbf}}, J_{\mathrm{a,sbf}}, J_{\mathrm{i,sbf}}) \cdot MBF_{\mathrm{T}}$ |
| $Mcm1 = G(\frac{k_{\mathrm{a,mcm}}}{c_{\mathrm{clb2}} \cdot mass} \cdot \mathrm{Clb2}, k_{\mathrm{i,mcm}}, J_{\mathrm{a,mcm}}, J_{\mathrm{i,mcm}}) \cdot Mcm1_{\mathrm{T}}$ |
| $Clb5_{\mathrm{T}} = Clb5 + C5 + C5\mathrm{P} + \mathrm{F5} + F5\mathrm{P}$ |
| $Clb2_{\mathrm{T}} = Clb2 + C2 + C2\mathrm{P} + F2 + F2\mathrm{P}$ |
| $Sic1_{\mathrm{T}} = Sic1 + Sic1\mathrm{P} + C2 + C2\mathrm{P} + C5 + C5\mathrm{P}$ |
| $Cdc6_{\mathrm{T}} = Cdc6 + Cdc6\mathrm{P} + F2 + F2\mathrm{P} + F5 + F5\mathrm{P}$ |
| $CKI_{\mathrm{T}} = Sic1_{\mathrm{T}} + Cdc6_{\mathrm{T}}$ |
| $RENT\mathrm{P} = Cdc14_{\mathrm{T}} - RENT - Cdc14$ |
| $Net1\mathrm{P} = Net1_{\mathrm{T}} - Net1 - Cdc14_{\mathrm{T}} + Cdc14$ |
| *(continued)* |

Supplementary Table S2. (continued)

| |
|---|
| $\mathrm{PE} = Esp1_{\mathrm{T}} - Esp1$ |
| $V_{\mathrm{d,b5}} = k'_{\mathrm{d,b5}} + k''_{\mathrm{d,b5}} \cdot Cdc20\mathrm{A}$ |
| $V_{\mathrm{d,b2}} = k'_{\mathrm{d,b2}} + k''_{\mathrm{d,b2}} \cdot Cdh1 + k_{\mathrm{d,b2p}} \cdot Cdc20\mathrm{A}$ |
| $V_{\mathrm{a,sbf}} = k_{\mathrm{a,sbf}} \cdot (e_{\mathrm{sbf,n2}} \cdot Cln2 + e_{\mathrm{sbf,n3}} \cdot (Cln3 + Bck2) + e_{\mathrm{sbf,b5}} \cdot Clb5)$ |
| $V_{\mathrm{i,sbf}} = k'_{\mathrm{i,sbf}} + k''_{\mathrm{i,sbf}} Clb2$ |
| $V_{\mathrm{kp,c1}} = k_{\mathrm{d1,c1}} + \frac{k_{\mathrm{d2,c1}}}{J_{\mathrm{d2,c1}} + \mathrm{Sic1T}} \cdot (e_{\mathrm{c1,n3}} \cdot Cln3 + e_{\mathrm{c1,k2}} \cdot Bck2 + e_{\mathrm{c1,n2}} \cdot Cln2 + e_{\mathrm{c1,b5}} \cdot Clb5 + e_{\mathrm{c1,b2}} \cdot Clb2)$ |
| $V_{\mathrm{kp,f6}} = k_{\mathrm{d1,f6}} + \frac{k_{\mathrm{d2,f6}}}{J_{\mathrm{d2,f6}} + Cdc6} \cdot (e_{\mathrm{f6,n3}} \cdot Cln3 + e_{\mathrm{f6,k2}} \cdot Bck2 + e_{\mathrm{f6,n2}} \cdot Cln2 + e_{\mathrm{f6,b5}} \cdot Clb5 + e_{\mathrm{f6,b2}} \cdot Clb2)$ |
| $V_{\mathrm{a,cdh}} = k'_{\mathrm{a,cdh}} + k''_{\mathrm{a,cdh}} \cdot Cdc14$ |
| $V_{\mathrm{i,cdh}} = k'_{\mathrm{i,cdh}} + k''_{\mathrm{i,cdh}} \cdot (e_{\mathrm{cdh,n3}} \cdot Cln3 + e_{\mathrm{cdh,n2}} \cdot Cln2 + e_{\mathrm{cdh,b2}} \cdot Clb2 + e_{\mathrm{cdh,b5}} \cdot Clb5)$ |
| $V_{\mathrm{pp,net}} = k'_{\mathrm{pp,net}} + k''_{\mathrm{pp,net}} \cdot Cdc55$ |
| $V_{\mathrm{kp,net}} = (k'_{\mathrm{kp,net}} + k''_{\mathrm{kp,net}} \cdot Cdc55) \cdot mass$ |
| $V_{\mathrm{d,55}} = k'_{\mathrm{d,55}} + k''_{\mathrm{d,55}} (J_{20,55} + Cdc20\mathrm{A}) \cdot \frac{J_{\mathrm{pds}}}{J_{\mathrm{pds}} + Pds1}$ |
| $V_{\mathrm{d,pds}} = k'_{\mathrm{d1,pds}} + k''_{\mathrm{d2,pds}} \cdot Cdc20\mathrm{A} + k''_{\mathrm{d3,pds}} \cdot Cdh1$ |

Reset rules: When the normalized concentration of Clb2 (denoted as $[\mathrm{Clb2}]_{\mathrm{n}}$) drops below $K_{\mathrm{ez}}$, we reset the auxiliary proteins $[\mathrm{BUD}]_{\mathrm{n}}$ and $[\mathrm{SPN}]_{\mathrm{n}}$ to zero, and divide all species in the cell, except for Cln3 and Bck2, between daughter and mother cells with a 40:60 ratio, according to observations by Di Talia *et al.* [1]. This ratio for Cln3 and Bck2 is set to 20:80 to match with the experimental observations in [2, 3]. When [Clb2]+[Clb5] drops below $K_{\mathrm{ez2}}$, $[\mathrm{ORI}]_{\mathrm{n}}$ is reset to zero. These auxiliary proteins are used as flags to specify particular events (check points in the budding yeast cell cycle). That is $[\mathrm{BUD}]_{\mathrm{n}} = 1$ indicates bud emergence, $[\mathrm{ORI}]_{\mathrm{n}} = 1$ specifies initiation of DNA synthesis and $[\mathrm{SPN}]_{\mathrm{n}} = 1$ signals that the chromosome alignment on spindle is completed.

The reset rules in our hybrid model are similar to the deterministic model by Chen *et al.* [4]. However, to prevent events from misfiring multiple times due to stochastic fluctuations in the hybrid model, we follow specific tactics. In our model, there are two types of events: *First*, events that are associated with the states of certain species that change across predefined thresholds in only one direction (increasing or decreasing), taking BUD as an example. *Second*, events that are associated with the states of certain species that change across predefined thresholds in both directions (increasing and decreasing), taking Clb2 as an instance.

For the first type of events, the thresholds are set to the original predefined values at the beginning of the cell cycle. Once an event is triggered, the corresponding threshold is increased by multiplying it with a large number (e.g., 1000) to prevent misfiring.

For the second type of events, we add two dummy events to prevent misfiring. Supplementary figure S1 illustrates an example which describes the procedure we use to monitor type-two events for Clb2. The goal is to monitor the occurrence of events 2 and 4. We add events 1 and 3 to prevent misfiring of events 2 and 4 in our hybrid stochastic model. At the beginning of cell cycle, all thresholds are set to very high values except for the first event which is set to $0.5K_{\text{ez}}$. Once event $i$ occurs, the threshold for $i$ is set to a very large value and the threshold for event $i+1$ is reset to its original predefined value. Using this procedure, we guarantee that events 2 and 4 take place in a correct manner in the presence of stochastic fluctuations.

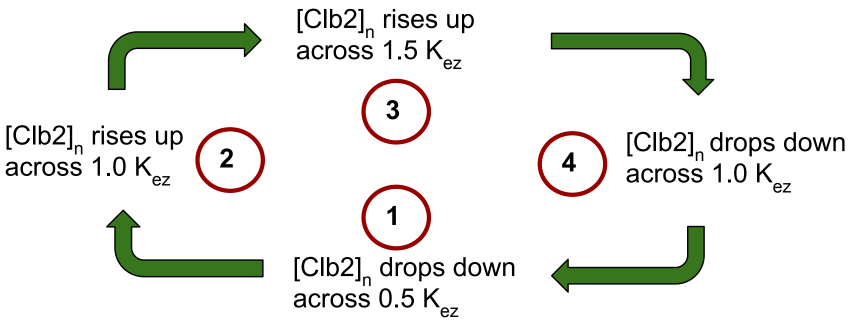


Supplementary Figure S 1: **Type-two event monitoring procedure [5].** Events 2 and 4 are the original events from the deterministic model [4]. Events 3 and 4 are the dummy events added to avoid misfiring of the original events in the presence of stochastic fluctuations. $K_{\text{ez}}$ is a dimensionless concentration.

Supplementary Table S3. Reaction channels and corresponding propensity functions for the slow subset in the hybrid model.

| Reaction | Propensity Function | Reaction | Propensity Function |
|---|---|---|---|
| $\phi \to mCdh1$ | $k_{\mathrm{s,mcdh1}}$ | $mCdh1 \to \phi$ | $k_{\mathrm{d,mcdh1}} \cdot mCdh1$ |
| $\phi \to mTem1$ | $k_{\mathrm{s,mtem1}}$ | $mTem1 \to \phi$ | $k_{\mathrm{d,mtem1}} \cdot mTem1$ |
| $\phi \to mCdc15$ | $k_{\mathrm{s,mcdc15}}$ | $mCdc15 \to \phi$ | $k_{\mathrm{d,mcdc15}} \cdot mCdc15$ |
| $\phi \to mCdc14$ | $k_{\mathrm{s,mcdc14}}$ | $mCdc14 \to \phi$ | $k_{\mathrm{d,mcdc14}} \cdot mCdc14$ |
| $\phi \to mNet1$ | $k_{\mathrm{s,mnet1}}$ | $mNet1 \to \phi$ | $k_{\mathrm{d,mnet1}} \cdot mNet1$ |
| $\phi \to mCdc55$ | $k_{\mathrm{s,mcdc55}}$ | $mCdc55 \to \phi$ | $k_{\mathrm{d,mcdc55}} \cdot mCdc55$ |
| $\phi \to mEsp1$ | $k_{\mathrm{s,mesp1}}$ | $mEsp1 \to \phi$ | $k_{\mathrm{d,mesp1}} \cdot mEsp1$ |
| $\phi \to mSBF$ | $k_{\mathrm{s,msbf}}$ | $mSBF \to \phi$ | $k_{\mathrm{d,msbf}} \cdot mSBF$ |
| $\phi \to mMBF$ | $k_{\mathrm{s,mmbf}}$ | $mMBF \to \phi$ | $k_{\mathrm{d,mmbf}} \cdot mMBF$ |
| $\phi \to mMcm1$ | $k_{\mathrm{s,mmcm1}}$ | $mMcm1 \to \phi$ | $k_{\mathrm{d,mmcm1}} \cdot mMcm1$ |
| $\phi \to mAPC$ | $k_{\mathrm{s,mapc}}$ | $mAPC \to \phi$ | $k_{\mathrm{d,mapc}} \cdot mAPC$ |
| $\phi \to mCln2$ | $k_{\mathrm{s,mcln2}} \cdot (k'_{\mathrm{s,n2}} \cdot c_{\mathrm{cln2}} \cdot mass + k''_{\mathrm{s,n2}} \cdot \frac{c_{\mathrm{cln2}}}{c_{\mathrm{sbf}}} \cdot SBF)$ | $mCln2 \to \phi$ | $k_{\mathrm{d,mcln2}} \cdot mCln2$ |
| $\phi \to mClb5$ | $k_{\mathrm{s,mclb5}} \cdot (k'_{\mathrm{s,b5}} \cdot c_{\mathrm{clb5}} \cdot mass + k''_{\mathrm{s,b5}} \cdot \frac{c_{\mathrm{clb5}}}{c_{\mathrm{mbf}}} \cdot MBF)$ | $mClb5 \to \phi$ | $k_{\mathrm{d,mclb5}} \cdot mClb5$ |
| $\phi \to mClb2$ | $k_{\mathrm{s,mclb2}} \cdot (k'_{\mathrm{s,b2}} \cdot c_{\mathrm{clb2}} \cdot mass + k''_{\mathrm{s,b2}} \cdot \frac{c_{\mathrm{clb2}}}{c_{\mathrm{mcm1}}} \cdot Mcm1)$ | $mClb2 \to \phi$ | $k_{\mathrm{d,mclb2}} \cdot mClb2$ |
| $\phi \to mSic1$ | $k_{\mathrm{s,msic1}} \cdot (k'_{\mathrm{s,c1}} \cdot c_{\mathrm{sic1}} \cdot mass + k''_{\mathrm{s,c1}} \frac{c_{\mathrm{sic1}}}{c_{\mathrm{swi5}}} \cdot Swi5)$ | $mSic1 \to \phi$ | $k_{\mathrm{d,msic1}} \cdot mSic1$ |
| $\phi \to mCdc6$ | $k_{\mathrm{s,mcdc6}} \cdot (k'_{\mathrm{s,f6}} \cdot c_{\mathrm{cdc6}} \cdot mass + k''_{\mathrm{s,f6}} \cdot \frac{c_{\mathrm{cdc6}}}{c_{\mathrm{swi5}}} \cdot Swi5 + k'''_{\mathrm{s,f6}} \cdot \frac{c_{\mathrm{cdc6}}}{c_{\mathrm{sbf}}} \cdot SBF)$ | $mCdc6 \to \phi$ | $k_{\mathrm{d,mcdc6}} \cdot mCdc6$ |

| | | | |
|---|---|---|---|
| $\phi \rightarrow mSwi5$ | $k_{\text{s,mswi5}} \cdot (k'_{\text{s,swi}} \cdot c_{\text{swi5}} \cdot mass + k''_{\text{s,swi}} \cdot \frac{c_{\text{swi5}}}{c_{\text{mcm1}}} \cdot Mcm1)$ | $mSwi5 \rightarrow \phi$ | $k_{\text{d,mswi5}} \cdot mSwi5$ |
| $\phi \rightarrow mCdc20$ | $k_{\text{s,mcdc20}} \cdot (k'_{\text{s,20}} \cdot c_{\text{cdc20}} \cdot mass + k''_{\text{s,20}} \cdot \frac{c_{\text{cdc20}}}{c_{\text{mcm1}}} \cdot Mcm1)$ | $mCdc20 \rightarrow \phi$ | $k_{\text{d,mcdc20}} \cdot mCdc20$ |
| $\phi \rightarrow mPds1$ | $k_{\text{s,mpds1}} \cdot (k'_{\text{s,pds}} \cdot c_{\text{pds1}} \cdot mass + k''_{\text{s1,pds}} \cdot \frac{c_{\text{pds1}}}{c_{\text{sbf}}} \cdot SBF + k''_{\text{s2,pds}} \cdot \frac{c_{\text{pds1}}}{c_{\text{mcm1}}} \cdot Mcm1)$ | $mPds1 \rightarrow \phi$ | $k_{\text{d,mpsd1}} \cdot mPds1$ |

Supplementary Table S4. Basal parameter values for the wild-type cell cycle. Parameters that start with lower case $k$ are rate constants ($\text{min}^{-1}$). All other parameter are dimensionless.

| | | | |
|---|---|---|---|
| $k_{\text{g}} = 0.0072$ | $k'_{\text{s,n2}} = 0$ | $k''_{\text{s,n2}} = 0.03$ | $k_{\text{d,n2}} = 0.12$ |
| $k'_{\text{s,b5}} = 0.0001$ | $k''_{\text{s,b5}} = 0.0007$ | $k'_{\text{d3,c1}} = 1$ | $k_{\text{di,b5}} = 0.06$ |
| $k_{\text{as,f5}} = 0.01$ | $k_{\text{d3,f6}} = 1$ | $k_{\text{di,f5}} = 0.01$ | $k_{\text{as,b5}} = 50$ |
| $k'_{\text{s,b2}} = 0.0003$ | $k''_{\text{s,b2}} = 0.0114$ | $k_{\text{di,b2}} = 0.05$ | $k_{\text{di,f2}} = 0.5$ |
| $k_{\text{as,b2}} = 50$ | $k_{\text{as,f2}} = 15$ | $k'_{\text{s,c1}} = 0.0036$ | $k''_{\text{s,c1}} = 0.0359$ |
| $k_{\text{pp,c1}} = 4$ | $k'_{\text{s,f6}} = 0.0059$ | $k''_{\text{s,f6}} = 0.0295$ | $k'''_{\text{s,f6}} = 0.0001$ |
| $k_{\text{pp,f6}} = 4$ | $k'_{\text{s,swi}} = 0.0007$ | $k''_{\text{s,swi}} = 0.0114$ | $k_{\text{d,swi}} = 0.08$ |
| $k_{\text{a,swi}} = 2$ | $k_{\text{i,swi}} = 0.05$ | $k_{\text{a,apc}} = 0.1$ | $k_{\text{i,apc}} = 0.15$ |
| $k'_{\text{s,20}} = 0.0014$ | $k''_{\text{s,20}} = 0.1364$ | $k'_{\text{a,20}} = 0.05$ | $k''_{\text{a,20}} = 0.2$ |
| $k_{\text{d,20}} = 0.3$ | $k_{\text{s,cdh}} = 0.0014$ | $k_{\text{d,cdh}} = 0.01$ | $k'_{\text{a,15}} = 0.002$ |
| $k''_{\text{a,15}} = 1$ | $k'''_{\text{a,15}} = 0.001$ | $k_{\text{i,15}} = 0.5$ | $k_{\text{s,14}} = 0.02$ |
| $k_{\text{d,14}} = 0.1$ | $k_{\text{d,net}} = 0.03$ | $k_{\text{di,rent}} = 1$ | $k_{\text{di,rentp}} = 2$ |
| $k_{\text{as,rent}} = 200$ | $k_{\text{as,rentp}} = 1$ | $k_{\text{s,net}} = 0.0134$ | $k_{\text{s,55}} = 0.0143$ |
| $k'_{\text{s,pds}} = 0$ | $k''_{\text{s1,pds}} = 0.0043$ | $k''_{\text{s2,pds}} = 0.0079$ | $k_{\text{di,esp}} = 0.5$ |
| $k_{\text{as,esp}} = 50$ | $k_{\text{s,ori}} = 2$ | $k_{\text{d,ori}} = 0.06$ | $k_{\text{s,bud}} = 0.2$ |
| $k_{\text{d,bud}} = 0.06$ | $k_{\text{s,spn}} = 0.1$ | $k_{\text{d,spn}} = 0.06$ | $k_{\text{s,tem1t}} = 0.0150$ |
| $k_{\text{d,tem1t}} = 0.046$ | $k_{\text{s,cdc15t}} = 0.0142$ | $k_{\text{d,cdc15t}} = 0.046$ | $k_{\text{s,esp1t}} = 0.0139$ |
| $k_{\text{d,esp1t}} = 0.046$ | $k_{\text{s,sbft}} = 0.007$ | $k_{\text{d,sbft}} = 0.0462$ | $k_{\text{a,mcm}} = 5$ |
| $k_{\text{s,mbft}} = 0.0066$ | $k_{\text{d,mbft}} = 0.046$ | $k_{\text{s,apct}} = 0.0065$ | $k_{\text{d,apct}} = 0.0462$ |
| $k_{\text{s,mcm1t}} = 0.0078$ | $k_{\text{d,mcm1t}} = 0.046$ | $k_{\text{i,mcm}} = 0.15$ | $k''_{\text{d3,pds}} = 0.04$ |
| $k_{\text{a,sbf}} = 0.38$ | $k'_{\text{i,sbf}} = 0.6$ | $k''_{\text{i,sbf}} = 8$ | $k'_{\text{d,b5}} = 0.01$ |
| $k'_{\text{d,b2}} = 0.003$ | $k''_{\text{d,b2}} = 0.4$ | $k_{\text{d,b2p}} = 0.15$ | $k_{\text{d1,c1}} = 0.01$ |
| $k_{\text{d2,c1}} = 1$ | $k''_{\text{d,b5}} = 0.16$ | $k_{\text{d1,f6}} = 0.01$ | $k_{\text{d2,f6}} = 1$ |
| $k'_{\text{a,cdh}} = 0.01$ | $k''_{\text{a,cdh}} = 0.8$ | $k'_{\text{i,cdh}} = 0.001$ | $k''_{\text{i,cdh}} = 0.08$ |
| $k'_{\text{pp,net}} = 0.05$ | $k''_{\text{pp,net}} = 3$ | $k'_{\text{kp,net}} = 0.01$ | $k''_{\text{kp,net}} = 0.6$ |
| $k'_{\text{d,55}} = 0.17$ | $k''_{\text{d,55}} = 2$ | $k'_{\text{d1,pds}} = 0.01$ | $k''_{\text{d2,pds}} = 0.2$ |

| Supplementary Table S4.(continued) | | | |
|---|---|---|---|
| $k_{\mathrm{s,n3}} = 0.0080$ | $k_{\mathrm{d,n3}} = 0.12$ | $k_{\mathrm{s,k2}} = 0.0048$ | $k_{\mathrm{d,k2}} = 0.12$ |
| $k_{\mathrm{mad2}} = 8$ for $[\mathrm{ORI}]_{\mathrm{n}} > 1$and $[\mathrm{SPN}]_{\mathrm{n}} < 1;$ otherwise 0.01 | | | |
| $k_{\mathrm{bud2}} = 1$ for $[\mathrm{ORI}]_{\mathrm{n}} > 1$ and $[\mathrm{SPN}]_{\mathrm{n}} < 1;$ otherwise 0.2 | | | |
| $k_{\mathrm{lte1}} = 1$ for $[\mathrm{SPN}]_{\mathrm{n}} > 1$ and $[\mathrm{Clb2}]_{\mathrm{n}} > 1;$ otherwise 0.1 | | | |
| $K_{\mathrm{ez}} = 0.3$ | $K_{\mathrm{ez2}} = 0.2$ | $J_{\mathrm{i,cdh}} = 0.03$ | $J_{\mathrm{a,apc}} = 0.1$ |
| $J_{\mathrm{i,apc}} = 0.1$ | $J_{\mathrm{a,cdh}} = 0.03$ | $J_{\mathrm{pds}} = 0.04$ | $J_{\mathrm{a,sbf}} = 0.01$ |
| $J_{\mathrm{n3}} = 6$ | $J_{\mathrm{a,tem}} = 0.1$ | $J_{\mathrm{i,tem}} = 0.1$ | $J_{\mathrm{spn}} = 0.14$ |
| $J_{\mathrm{d2,f6}} = 0.05$ | $J_{\mathrm{a,mcm}} = 0.1$ | $J_{\mathrm{i,mcm}} = 0.1$ | $J_{\mathrm{d2,c1}} = 0.05$ |
| $J_{\mathrm{i,sbf}} = 0.01$ | $J_{20,55} = 0.15$ | $e_{\mathrm{cdh,b2}} = 1.2$ | $e_{\mathrm{cdh,b5}} = 8$ |
| $e_{\mathrm{bud,n3}} = 0.05$ | $e_{\mathrm{ori,b5}} = 0.9$ | $e_{\mathrm{ori,b2}} = 0.45$ | $e_{\mathrm{bud,n2}} = 0.12$ |
| $e_{\mathrm{sbf,b5}} = 2$ | $e_{\mathrm{bud,b5}} = 0.25$ | $e_{\mathrm{sbf,n2}} = 2$ | $e_{\mathrm{sbf,n3}} = 10$ |
| $e_{\mathrm{c1,b5}} = 0.1$ | $e_{\mathrm{c1,n3}} = 0.3$ | $e_{\mathrm{c1,k2}} = 0.03$ | $e_{\mathrm{c1,n2}} = 0.06$ |
| $e_{\mathrm{f6,n2}} = 0.06$ | $e_{\mathrm{c1,b2}} = 0.45$ | $e_{\mathrm{f6,n3}} = 0.3$ | $e_{\mathrm{f6,k2}} = 0.03$ |
| $e_{\mathrm{cdh,n2}} = 0.4$ | $e_{\mathrm{f6,b5}} = 0.1$ | $e_{\mathrm{f6,b2}} = 0.55$ | $e_{\mathrm{cdh,n3}} = 0.25$ |
| $c_{\mathrm{clb2}} = 479.2$ | $c_{\mathrm{cln2}} = 239.6$ | $c_{\mathrm{cln3}} = 599$ | $c_{\mathrm{bck2}} = 599$ |
| $c_{\mathrm{cdh1}} = 1198$ | $c_{\mathrm{ori}} = 119.8$ | $c_{\mathrm{bud}} = 119.8$ | $c_{\mathrm{spn}} = 119.8$ |
| $c_{\mathrm{tem1}} = 178.2$ | $c_{\mathrm{clb5}} = 479.2$ | $c_{\mathrm{cdc20}} = 1797$ | $c_{\mathrm{sic1}} = 479.2$ |
| $c_{\mathrm{cdc6}} = 479.2$ | $c_{\mathrm{cdc14}} = 275.54$ | $c_{\mathrm{net1}} = 275.54$ | $c_{\mathrm{cdc15}} = 479.2$ |
| $c_{\mathrm{pds1}} = 39.5$ | $c_{\mathrm{esp1}} = 39.5$ | $c_{\mathrm{sbf}} = 479.2$ | $c_{\mathrm{mbf}} = 479.2$ |
| $c_{\mathrm{apc}} = 263.56$ | $c_{\mathrm{cdc55}} = 4193$ | $c_{\mathrm{rent}} = 4193$ | $c_{\mathrm{mcm1}} = 4193$ |
| $k_{\mathrm{s,mCdc15}} = 0.45$ | $k_{\mathrm{d,mCdc15}} = 0.14$ | $k_{\mathrm{s,mTem1}} = 0.43$ | $k_{\mathrm{d,mTem1}} = 0.14$ |
| $k_{\mathrm{s,mNet1}} = 0.5$ | $k_{\mathrm{d,mNet1}} = 0.1$ | $k_{\mathrm{s,mCdc14}} = 1.5$ | $k_{\mathrm{d,mCdc14}} = 0.14$ |
| $k_{\mathrm{s,mEsp1}} = 0.3$ | $k_{\mathrm{d,mEsp1}} = 0.08$ | $k_{\mathrm{s,mCdc55}} = 0.97$ | $k_{\mathrm{d,mCdc55}} = 0.14$ |
| $k_{\mathrm{s,mMBF}} = 0.97$ | $k_{\mathrm{d,mMBF}} = 0.14$ | $k_{\mathrm{s,mSBF}} = 0.94$ | $k_{\mathrm{d,mSBF}} = 0.14$ |
| $k_{\mathrm{s,mPds1}} = 0.24$ | $k_{\mathrm{d,mPds1}} = 0.14$ | $k_{\mathrm{s,mMcm1}} = 0.8$ | $k_{\mathrm{d,mMcm1}} = 0.14$ |
| $k_{\mathrm{s,mAPC}} = 0.97$ | $k_{\mathrm{d,mAPC}} = 0.14$ | $k_{\mathrm{s,mCln2}} = 0.0117$ | $k_{\mathrm{d,mCln2}} = 0.23$ |
| $k_{\mathrm{s,mClb5}} = 0.3916$ | $k_{\mathrm{d,mClb5}} = 0.14$ | $k_{\mathrm{s,mClb2}} = 0.0034$ | $k_{\mathrm{d,mClb2}} = 0.35$ |

Supplementary Table S4.(continued)

| | | | |
|---|---|---|---|
| $k_{\mathrm{s,mSic1}} = 0.0067$ | $k_{\mathrm{d,mSic1}} = 0.14$ | $k_{\mathrm{s,mCdc6}} = 0.0083$ | $k_{\mathrm{d,mCdc6}} = 0.14$ |
| $k_{\mathrm{s,mSwi5}} = 0.0016$ | $k_{\mathrm{d,mSwi5}} = 0.14$ | $k_{\mathrm{s,mCdc20}} = 0.0001$ | $k_{\mathrm{d,mCdc20}} = 0.14$ |
| $k_{\mathrm{s,mCdh1}} = 0.97$ | $k_{\mathrm{d,mCdh1}} = 0.14$ | | |

The FORTRAN code that implements the model described in Supplementary Table S1-S4 is provided in Supplementary Code. We notice that the parameters listed in Supplementary Table S4 are the values that are computed after converting the concentration of species into population. In the code, however, to provide more flexibility in tuning the parameters, the original parameters from Chen's model are listed and then converted. See the README file that further elaborates on the Supplementary Code.

Supplementary Table S5: List of mutant strains. Simulation results of the hybrid stochastic model (in column 5) are compared with experimental data (in column 4. The parameter modification column reports the changes made to the parameter values in order to simulate each strain. 'V' means 'viable' and 'In' means inviable. We generate a sufficiently large population of mother and daughter cells, expanding over time. In column 5 the computed probability of division is reported for each mutant. Red typeface in column 5 indicates disagreement between the hybrid model and the experimental observations.

| No | Mutant strain | Parameter modification | Experiment | Hybrid model |
|---|---|---|---|---|
| 1 | Wild-type (WT) on glucose | - | V | V : 0.97 |
| 2.1 | WT on galactose | $k_g$=0.004621 (MDT = 150) | V | V : 0.92 |
| 2.2 | WT on raffinose | $k_g = 0.00433$ (MDT = 160) | V | V : 0.90 |
| 3 | *cln*1Δ *cln*2Δ | $k''_{s,n2} = 0$ | V | V : 0.96 |
| 4 | *GAL-CLN*2 *cln*1Δ *cln*2Δ | $k'_{s,n2} = 0.12$, $k''_{s,n2} = 0$, $k_g = 0.004621$ | V | V : 0.96 |
| 5 | *cln*1Δ *cln*2Δ *sic*1Δ | $k''_{s,n2} = 0$, $k'_{s,c1} = k''_{s,c1} = 0$ | V | V : 0.75 |
| 6 | *cln*1Δ *cln*2Δ *cdh*1Δ | $k''_{s,n2} = 0$, $k_{s,cdh} = 0$, init CDH1T = CDH1 = 0 | V | In : 0.27 |
| 7 | *GAL-CLN*2 *cln*1Δ *cln*2Δ *cdh*1Δ | $k'_{s,n2} = 0.12$, $k''_{s,n2} = 0$, $k_{s,cdh} = 0$, $k_g = 0.004621$, init CDH1T = CDH1 = 0 | V | V : 0.76 |
| 8 | *cln*3Δ | $k_{s,n3} = 0$ | V | V : 0.98 |
| 9 | *GAL-CLN*3 | $k_{s,n3} = 5 \times k_{s,n3}$, $k_g = 0.004621$ | V | V : 0.94 |
| 10 | *bck*2Δ | $k_{s,k2} = 0$ | V | V : 0.97 |
| 11 | multi-copy *BCK*2 | $k_{s,k2} = 5 \times k_{s,k2}$ | V | V : 0.96 |
| 12 | *cln*1Δ *cln*2Δ *bck*2Δ | $k''_{s,n2} = 0$, $k_{s,k2} = 0$ | V | V : 0.84 |
| 13 | *cln*3Δ *bck*2Δ | $k_{s,n3} = 0$, $k_{s,k2} = 0$ | In | In : 0.10 |
| 14 | *cln*3Δ *bck*2Δ *GAL-CLN*2 *cln*1Δ *cln*2Δ | $k_{s,n3} = 0$, $k_{s,k2} = 0$, $k'_{s,n2} = 0.12$, $k''_{s,n2} = 0$, $k_g = 0.004621$ | V | V : 0.96 |

| | | | | |
|---|---|---|---|---|
| 15 | *cln*3Δ *bck*2Δ multi-copy *CLN*2 | $k_{\rm s,n3} = 0$, $k_{\rm s,k2} = 0$<br>$k''_{\rm s,n2} = 5\times k''_{\rm s,n2}$ | In | In : 0.40 |
| 16 | *cln*3Δ *bck*2Δ *sic*1Δ | $k_{\rm s,n3} = 0$, $k_{\rm s,k2} = 0$,<br>$k'_{\rm s,c1} = k''_{\rm s,c1} = 0$ | In | In : 0.50 |
| 17 | *cln*1Δ *cln*2Δ *cln*3Δ | $k'_{\rm s,n2} = k''_{\rm s,n2} = 0$, $k_{\rm s,n3} = 0$ | In | In : 0.41 |
| 18 | *cln*1Δ *cln*2Δ *cln*3Δ *GAL-CLN*2 | $k''_{\rm s,n2} = 0$, $k_{\rm s,n3} = 0$,<br>$k'_{\rm s,n2} = 0.12$, $k_{\rm g} = 0.004621$ | V | V : 0.96 |
| 19 | *cln*1Δ *cln*2Δ *cln*3Δ *GAL-CLN*3 | $k'_{\rm s,n2} = k''_{\rm s,n2} = 0$,<br>$k_{\rm s,n3} = 10\times k_{\rm s,n3}$ ,<br>$k_{\rm g} = 0.004621$ | V | V : 0.85 |
| 20 | *cln*1Δ *cln*2Δ *cln*3Δ *sic*1Δ | $k'_{\rm s,n2} = 0$, $k''_{\rm s,n2} = 0$,<br>$k_{\rm s,n3} = 0$, $k'_{\rm s,c1} = k''_{\rm s,c1} = 0$ | V | V : 0.76 |
| 21 | *cln*1Δ *cln*2Δ *cln*3Δ *cdh*1Δ | $k'_{\rm s,n2} = k''_{\rm s,n2} = 0$,<br>$k_{\rm s,n3} = 0$, $k_{\rm s,cdh} = 0$,<br>init CDH1 = CDH1T = 0 | In | In : 0.03 |
| 22 | *cln*1Δ *cln*2Δ *cln*3Δ multi-copy *CLB*5 | $k'_{\rm s,n2} = k''_{\rm s,n2} = 0$,<br>$k_{\rm s,n3} = 0$, $k'_{\rm s,b5} = 2\times k'_{\rm s,b5}$,<br>$k''_{\rm s,b5} = 2\times k''_{\rm s,b5}$ | V | V : 0.88 |
| 23 | *cln*1Δ *cln*2Δ *cln*3Δ *GAL-CLB*5 | $k'_{\rm s,n2} = 0$, $k''_{\rm s,n2} = 0$,<br>$k_{\rm s,n3} = 0$, $k'_{\rm s,b5} = 15\times k'_{\rm s,b5}$,<br>$k_{\rm g} = 0.004621$ | V | V : 0.99 |
| 24 | *cln*1Δ *cln*2Δ *cln*3Δ multi-copy *BCK*2 | $k'_{\rm s,n2} = k''_{\rm s,n2} = 0$,<br>$k_{\rm s,n3} = 0$, $k_{\rm s,k2} = 5\times k_{\rm s,k2}$ | V | V : 0.84 |
| 25 | *cln*1Δ *cln*2Δ *cln*3Δ *GAL-CLB*2 | $k'_{\rm s,n2} = k''_{\rm s,n2} = 0$,<br>$k_{\rm s,n3} = 0$, $k'_{\rm s,b2} = 2\times k'_{\rm s,b2}$,<br>$k_{\rm g} = 0.004621$ | In | In : 0.10 |

| | | | | |
|---|---|---|---|---|
| 26 | *cln*1Δ *cln*2Δ *cln*3Δ *apc-ts* | $k'_{\rm s,n2} = k''_{\rm s,n2} = 0$, $k_{\rm s,n3} = 0$, $k_{\rm s,cdh} = 0$, $k'_{\rm s,20} = k''_{\rm s,20} = 0$, init CDH1 = CDH1T = 0 | In | In : 0 |
| 27 | *sic*1Δ | $k'_{\rm s,c1} = k''_{\rm s,c1} = 0$ | V | V : 0.87 |
| 28 | *GAL-SIC*1 | $k'_{\rm s,c1} = 10\times k'_{\rm s,c1}$, $k_{\rm g} = 0.004621$ | V | V : 0.94 |
| 29 | *GAL-SIC*1*db*Δ | $k'_{\rm s,c1} = 10\times k'_{\rm s,c1}$, $k_{\rm d3,c1} = 0$, $k_{\rm g} = 0.004621$ | In | In : 0 |
| 30 | *GAL-SIC*1 *cln*1Δ *cln*2Δ | $k'_{\rm s,c1} = 10\times k'_{\rm s,c1}$, $k''_{\rm s,n2} = 0$, $k_{\rm g} = 0.004621$ | In | In : 0 |
| 31 | *GAL-SIC*1 *GAL-CLN*2 *cln*1Δ *cln*2Δ | $k'_{\rm s,c1} = 10\times k'_{\rm s,c1}$, $k'_{\rm s,n2} = 0.12$, $k''_{\rm s,n2} = 0$, $k_{\rm g} = 0.004621$ | V | V : 0.98 |
| 32 | *GAL-SIC*1 *cln*1Δ *cln*2Δ *cdh*1Δ | $k'_{\rm s,c1} = k''_{\rm s,c1} = 0.12$, $k''_{\rm s,n2} = k_{\rm s,cdh} = 0$, $k_{\rm g} = 0.004621$, init CDH1T = CDH1 = 0 | In | In : 0 |
| 33 | *GAL-SIC*1 *GAL-CLN*2 *cln*1Δ *cln*2Δ *cdh*1Δ | $k'_{\rm s,c1} = 0.12$, $k'_{\rm s,n2} = 0.12$, $k''_{\rm s,n2} = 0$, $k_{\rm s,cdh} = 0$, $k_{\rm g} = 0.004621$, init CDH1T = CDH1 = 0 | V | V : 0.93 |
| 34 | *cdh*1Δ | $k_{\rm s,cdh} = 0$, init CDH1T = CDH1 = 0 | V | In : 0.44 |
| 35 | *Cdh*1 constitutively active | $k''_{\rm i,cdh} = 0$, $k_{\rm g} = 0.004621$, $k_{\rm s,cdh} = 3\times k_{\rm s,cdh}$ | In | In : 0.34 |

| | | | | |
|---|---|---|---|---|
| 36 | *sic*1Δ *cdh*1Δ | $k'_{\mathrm{s,c1}} = k''_{\mathrm{s,c1}} = 0$, $k_{\mathrm{s,cdh}} = 0$, init CDH1T = CDH1 = 0 | In | In : 0 |
| 37 | *sic*1Δ *cdh*1Δ *GAL-CDC*20 | $k'_{\mathrm{s,c1}} = k''_{\mathrm{s,c1}} = 0$, $k_{\mathrm{s,cdh}} = 0$, $k'_{\mathrm{s,20}} = 10$, $k_{\mathrm{g}} = 0.004621$ | V | partially viable: 0.71 |
| 38 | *cdc*6Δ2-49 | $k'_{\mathrm{s,f6}} = k''_{\mathrm{s,f6}} = k'''_{\mathrm{s,f6}} = 0$ | V | V : 0.97 |
| 39 | *sic*1Δ *cdc*6Δ2-49 | $k'_{\mathrm{s,c1}} = k''_{\mathrm{s,c1}} = 0$, $k'_{\mathrm{s,f6}} = k''_{\mathrm{s,f6}} = k'''_{\mathrm{s,f6}} = 0$ | V | In : 0.08 |
| 40 | *cdh*1Δ *cdc*6Δ2-49 | $k'_{\mathrm{s,f6}} = k''_{\mathrm{s,f6}} = k'''_{\mathrm{s,f6}} = 0$, $k_{\mathrm{s,cdh}} = 0$, init CDH1T = CDH1 = 0 | V | In : 0.16 |
| 41[1] | *sic*1Δ *cdc*6Δ2-49 *cdh*1Δ | $k'_{\mathrm{s,c1}} = k''_{\mathrm{s,c1}} = 0$, $k_{\mathrm{s,cdh}} = 0$, $k'_{\mathrm{s,f6}} = k''_{\mathrm{s,f6}} = k'''_{\mathrm{s,f6}} = 0$, init CDH1T = CDH1 = 0 | In | In : 0 |
| 42 | *sic*1Δ *cdc*6Δ2-49 *cdh*1Δ *GAL-CDC*20 | $k'_{\mathrm{s,c1}} = k''_{\mathrm{s,c1}} = 0$, $k_{\mathrm{s,cdh}} = 0$, $k'_{\mathrm{s,f6}} = k''_{\mathrm{s,f6}} = k'''_{\mathrm{s,f6}} = 0$, $k'_{\mathrm{s,20}} = 4$, $k_{\mathrm{g}} = 0.004621$, init CDH1T = CDH1 = 0 | V | In : 0.19 |
| 43 | *swi*5Δ | $k'_{\mathrm{s,swi}} = k''_{\mathrm{s,swi}} = 0$ | V | In : 0.62 |
| 44 | *swi*5Δ *GAL-CLB*2 | $k'_{\mathrm{s,swi}} = k''_{\mathrm{s,swi}} = 0$, $k_{\mathrm{s,b2}} = 0.12$, $k_{\mathrm{g}} = 0.004621$ | In | In : 0.34 |
| 45[1] | *swi*5Δ *cdh*1Δ | $k'_{\mathrm{s,swi}} = k''_{\mathrm{s,swi}} = 0$, $k_{\mathrm{s,cdh}} = 0$, init CDH1T = CDH1 = 0 | In | In : 0 |

[1] Experimental observations show that *sic*1Δ *cdc*6Δ2-49 *cdh*1Δ (mutant # 41) and *swi*5Δ *cdh*1Δ (mutant # 45) strains are not able to undergo the cell division and thus are inviable [6]; both strains have similar phenotype and become arrested as binucleate cells with 4C DNA content. Our simulation (similar to Chen's deterministic simulation [4]) shows that the cells are inviable, however, they become arrested in telophase with 2C DNA content. We consider this simulation result as an inconsistency with experimental observations.

| | | | | |
|---|---|---|---|---|
| 46 | *swi*5Δ *cdh*1Δ *GAL-SIC*1 | $k'_{\rm s,swi}$= $k''_{\rm s,swi}$= 0, $k_{\rm s,cdh}$= 0, $k'_{\rm s,c1}$ = 10×$k'_{\rm s,c1}$, $k_{\rm g}$ = 0.004621, init CDH1T = CDH1 = 0 | V | V : 0.83 |
| 47 | *clb*1Δ *clb*2Δ | $k'_{\rm s,b2} = k''_{\rm s,b2} = 0$ | In | In : 0 |
| 48 | *CLB*1 *clb*2Δ | $k'_{\rm s,b2}$ = 0.0003, $k''_{\rm s,b2}$ = 0.013 (33% of WT) | V | V : 0.83 |
| 49 | *GAL-CLB*2 | $k'_{\rm s,b2}$ = 0.12, $k_{\rm g}$ = 0.004621 | V | V : 0.87 |
| 50 | multi-copy *GAL-CLB*2 | $k'_{\rm s,b2}$ = 0.96 (8 copies of *GAL-CLB*2), $k_{\rm g}$ = 0.004621 | In | In : 0 |
| 51[2] | *CLB*1 *clb*2Δ *cdh*1Δ | $k'_{\rm s,b2}$ = 0.0003, $k''_{\rm s,b2}$ = 0.013 (33% of WT), $k_{\rm s,cdh}$ = 0, init CDH1T = CDH1 = 0 | ga : V gl : V? | ga :V: 0.79, gl : In : 0.62 |
| 52[3] | *CLB*1 *clb*2Δ *pds*1Δ | $k'_{\rm s,b2}$ = 0.0003, $k''_{\rm s,b2}$ = 0.013 (33% of WT), $k''_{\rm s1,pds} = k''_{\rm s2,pds} = 0$ | In | In |
| 53 | *GAL-CLB*2 *sic*1Δ | $k'_{\rm s,b2}$ = 0.12, $k'_{\rm s,c1} = k''_{\rm s,c1}$ = 0, $k_{\rm g}$ = 0.004621 | In | In : 0.14 |

[2]Experimental observations show that *CLB*1 *clb*2Δ *cdh*1Δ grows well on galactose, however, exhibits poor viability on glucose [7]. That is why on column 4 the viability of the mutant is followed by a question mark. 'gl' refers to growth in glucose medium; 'ga' to galactose.

[3]Simulation results show that *CLB*1 *clb*2Δ *pds*1Δ (mutant # 52), and *pds*1Δ (mutant # 80) strains are able to exit mitosis, however, Esp1 is active throughout the cell cycle (even before spindle alignment); thus, according to our viability rule these mutants are inviable.

| | | | | |
|---|---|---|---|---|
| 54 | *GAL-CLB2 cdh1Δ* | $k'_{\rm s,b2} = 0.12$, $k_{\rm s,cdh} = 0$, $k_{\rm g} = 0.004621$, init CDH1T = CDH1 = 0 | In | In : 0 |
| 55 | *CLB2-dbΔ* | $k_{\rm d,b2p} = 0$, $k''_{\rm d,b2} = 0.03$ (8.5% activity left due to KEN box). | In | In : 0.24 |
| 56 | *CLB2-dbΔ* on galactose | $k_{\rm d,b2p} = 0$, $k''_{\rm d,b2} = 0.03$, $k_{\rm g} = 0.004621$ | In | In : 0.59 |
| 57 | *CLB2-dbΔ* multi-copy *SIC1* | $k_{\rm d,b2p} = 0$, $k''_{\rm d,b2} = 0.03$, $k'_{\rm s,c1} = 10 \times k'_{\rm s,c1}$, $k''_{\rm s,c1} = 10 \times k''_{\rm s,c1}$ | V | V : 0.79 |
| 58 | *CLB2-dbΔ GAL-SIC1* | $k_{\rm d,b2p} = 0$, $k''_{\rm d,b2} = 0.03$, $k'_{\rm s,c1} = 10 \times k'_{\rm s,c1}$, $k_{\rm g} = 0.004621$ | V | V : 0.79 |
| 59 | *CLB2-dbΔ* multi-copy *CDC6* | $k_{\rm d,b2p} = 0$, $k''_{\rm d,b2} = 0.03$, $k'_{\rm s,f6} = 5 \times k'_{\rm s,f6}$, $k''_{\rm s,f6} = 5 \times k''_{\rm s,f6}$, $k'''_{\rm s,f6} = 5 \times k'''_{\rm s,f6}$ | V | In : 0.64 |
| 60 | *CLB2-dbΔ clb5Δ* | $k_{\rm d,b2p} = 0$, $k''_{\rm d,b2} = 0.03$, $k'_{\rm s,b5} = k''_{\rm s,b5} = 0$ | In | In : 0.24 |
| 61 | *CLB2-dbΔ clb5Δ* on raffinose | $k_{\rm d,b2p} = 0$, $k''_{\rm d,b2} = 0.03$, $k'_{\rm s,b5} = k''_{\rm s,b5} = 0$, $k_{\rm g} = 0.004621$ | partially viable | partially viable: 0.65 |
| 62 | *GAL-CLB2-dbΔ* | $k_{\rm d,b2p} = 0$, $k''_{\rm d,b2} = 0.03$, $k''_{\rm s,b2} = 0.12$, $k_{\rm g} = 0.004621$ | In | In : 0.03 |
| 63 | *clb5Δ clb6Δ* | $k'_{\rm s,b5} = k''_{\rm s,b5} = 0$ | V | V : 0.99 |

| | | | | |
|---|---|---|---|---|
| 64 | *cln*1Δ *cln*2Δ *clb*5Δ *clb*6Δ | $k'_{\mathrm{s,n2}} = k''_{\mathrm{s,n2}} = 0$, $k'_{\mathrm{s,b5}} = k''_{\mathrm{s,b5}} = 0$ | In | In : 0 |
| 65 | *GAL-CLB*5 | $k'_{\mathrm{s,b5}} = 15{\times}k'_{\mathrm{s,b5}}$, $k_{\mathrm{g}} = 0.004621$ | V | V : 0.94 |
| 66[4] | *GAL-CLB*5 *sic*1Δ | $k'_{\mathrm{s,b5}} = 15{\times}k'_{\mathrm{s,b5}}$, $k'_{\mathrm{s,c1}} = k''_{\mathrm{s,c1}} = 0$, $k_{\mathrm{g}} = 0.004621$ | In | In |
| 67 | *GAL-CLB*5 *cdh*1Δ | $k'_{\mathrm{s,b5}} = 10{\times}k'_{\mathrm{s,b5}}$, $k_{\mathrm{s,cdh}} = 0$, $k_{\mathrm{g}} = 0.004621$, init CDH1T = CDH1 = 0 | In | In : 0.59 |
| 68 | *CLB*5-*db*Δ | $k''_{\mathrm{d,b5}} = 0$ | V | V : 0.95 |
| 69[4] | *CLB*5-*db*Δ *sic*1Δ | $k''_{\mathrm{d,b5}} = 0$, $k'_{\mathrm{s,c1}} = k''_{\mathrm{s,c1}} = 0$ | In | In |
| 70 | *CLB*5-*db*Δ *pds*1Δ | $k''_{\mathrm{d,b5}} = 0$, $k''_{\mathrm{s1,pds}} = k''_{\mathrm{s2,pds}} = 0$ | V | V : 0.97 |
| 71 | *CLB*5-*db*Δ *pds*1Δ *cdc*20Δ | $k''_{\mathrm{d,b5}} = 0$, $k''_{\mathrm{s1,pds}} = k''_{\mathrm{s2,pds}} = 0$, $k'_{\mathrm{s,20}} = k''_{\mathrm{s,20}} = 0$ | In | partially viable : 0.68 |
| 72[4] | *GAL-CLB*5-*db*Δ | $k'_{\mathrm{s,b5}} = 15{\times}k'_{\mathrm{s,b5}}$, $k''_{\mathrm{d,b5}} = 0$, $k_{\mathrm{g}} = 0.004621$ | In | In |
| 73 | *cdc*20-*ts* | $k'_{\mathrm{s,20}} = k''_{\mathrm{s,20}} = 0$ | In | In : 0.31 |
| 74 | *cdc*20Δ *clb*5Δ | $k'_{\mathrm{s,20}} = k''_{\mathrm{s,20}} = 0$, $k'_{\mathrm{s,b5}} = k''_{\mathrm{d,b5}} = 0$ | In | In : 0.58 |
| 75 | *cdc*20Δ *pds*1Δ | $k'_{\mathrm{s,20}} = k''_{\mathrm{s,20}} = 0$, $k''_{\mathrm{s1,pds}} = k''_{\mathrm{s2,pds}} = 0$ | In | partially viable : 0.68 |

[4]Simulation results show that mutant strains *GAL-CLB*5 *sic*1Δ (mutant # 66), *CLB*5-*db*Δ *sic*1Δ (mutant # 69), and *GAL-CLB*5-*db*Δ (mutant # 72) are able to exit mitosis, however, [ORI] is not relicensed (due to high activity of Clb5 in G1 phase); thus, according to our viability rule, these mutants are inviable.

| | | | | |
|---|---|---|---|---|
| 76 | *cdc*20Δ *pds*1Δ *clb*5Δ | $k'_{\mathrm{s,20}} = k''_{\mathrm{s,20}} = 0$, $k'_{\mathrm{s,b5}} = k''_{\mathrm{d,b5}} = 0$, $k''_{\mathrm{s1,pds}} = k''_{\mathrm{s2,pds}} = 0$ | V | V : 0.95 |
| 77 | *GAL-CDC*20 | $k'_{\mathrm{s,20}} = 6$, $k_{\mathrm{g}} = 0.004621$ | In | V : 0.99 |
| 78 | *cdc*20-*ts mad*2Δ | $k'_{\mathrm{s,20}} = k''_{\mathrm{s,20}} = 0$, $k_{\mathrm{mad2}} = 0.01$ | In | In : 0.30 |
| 79 | *cdc*20-*ts bub*2Δ | $k'_{\mathrm{s,20}} = k''_{\mathrm{s,20}} = 0$, $k_{\mathrm{bub2}} = 0$, init BUB2 = 0 | In | In : 0.24 |
| 80[3] | *pds*1Δ | $k''_{\mathrm{s1,pds}} = k''_{\mathrm{s2,pds}} = 0$ | In | In |
| 81[5] | *esp*1-*ts* | $k_{\mathrm{as,esp}} = 0.1$, $k_{\mathrm{di,esp}} = 0.002$, $(1/500 \times \mathrm{WT})$ | In | In |
| 82[6] | *PDS*1-*db*Δ | $k''_{\mathrm{d2,pds}} = k''_{\mathrm{d3,pds}} = 0$ | In | In |
| 83[6] | *GAL-PDS*1-*db*Δ | $k'_{\mathrm{s,pds}} = 0.1$, $k_{\mathrm{g}} = 0.004621$ $k''_{\mathrm{d2,pds}} = k''_{\mathrm{d3,pds}} = 0$ | In | In |
| 84[6] | *GAL-PDS*1-*db*Δ *esp*1-*ts* | $k'_{\mathrm{s,pds}} = 0.1$, $k''_{\mathrm{d2,pds}} = k''_{\mathrm{d3,pds}} = 0$, $k_{\mathrm{as,esp}} = 0.1$, $k_{\mathrm{di,esp}} = 0.002$, $(1/500 \times \mathrm{WT})$, $k_{\mathrm{g}} = 0.004621$ | In | In |
| 85 | *GAL-ESP*1 *cdc*20-*ts* | $k'_{\mathrm{s,20}} = k''_{\mathrm{s,20}} = 0$, $k'_{\mathrm{s,esp1t}} = 5 \times k'_{\mathrm{s,esp1t}}$, $k_{\mathrm{g}} = 0.004621$ | In | In : 0.48 |
| 86 | *tem*1Δ | $k''_{\mathrm{a,15}} = 0.002$ $(= k'_{\mathrm{a,15}})$, $k_{\mathrm{s,mTem1}} = 0$ | In | V : 0.76 |

[5] Simulation results show that *esp*1-*ts* strains are able to exit mitosis while the sister chromatids are not yet separated. Therefore, according to our viability rule this mutant is inviable.

[6] Simulation results show that *PDS*1-*db*Δ (mutant # 82), *GAL-PDS*1-*db*Δ (mutant # 83), and *GAL-PDS*1-*db*Δ *esp*1-*ts* (mutant # 84) strains are able to exit mitosis, however, Esp1 is inactive throughout the cell cycle which means sister chromatids are still attached. Therefore, according to our viability rule these mutants are inviable.

| | | | | |
|---|---|---|---|---|
| 87 | *GAL-TEM1* | $k_{\mathrm{s,tem1t}} = 5{\times}k_{\mathrm{s,tem1t}}$,<br>$k_{\mathrm{g}} = 0.004621$ | V | V : 0.91 |
| 88 | *tem1-ts* multi-copy *CDC15* | $k''_{\mathrm{a,15}} = 0.002$,<br>$k_{\mathrm{s,mTem1}} = 0$, $k_{\mathrm{s,tem1t}} = 0$,<br>$k_{\mathrm{s,cdc15t}} = 5{\times}k_{\mathrm{s,cdc15t}}$ | V | V : 0.77 |
| 89 | *tem1-ts GAL-CDC15* | $k''_{\mathrm{a,15}} = 0.002$,<br>$k_{\mathrm{s,mTem1}} = 0$, $k_{\mathrm{s,tem1t}} = 0$,<br>$k_{\mathrm{s,cdc15t}} = 15{\times}k_{\mathrm{s,cdc15t}}$,<br>$k_{\mathrm{g}} = 0.004621$ | V | V : 0.81 |
| 90 | *tem1Δ net1-ts* | $k''_{\mathrm{a,15}} = 0.002$,<br>$k_{\mathrm{s,mTem1}} = 0$, $k_{\mathrm{s,tem1t}} = 0$,<br>$k_{\mathrm{as,rent}}=10$, $k_{\mathrm{as,rentp}}=0.05$,<br>(0.05× WT) | V | V : 0.99 |
| 91 | *tem1-ts* multi-copy *CDC14* | $k''_{\mathrm{a,15}} = 0.002$,<br>$k_{\mathrm{s,14}} = 0.4$ (2 copies),<br>$k_{\mathrm{s,mTem1}} = k_{\mathrm{s,tem1t}} = 0$ | V | V : 0.96 |
| 92 | *cdc15Δ* | $k''_{\mathrm{kp,net}} = 0$, $k_{\mathrm{s,cdc15t}} = 0$ | In | V : 0.77 |
| 93 | multi-copy *CDC15* | $k_{\mathrm{s,cdc15t}} = 5{\times}k_{\mathrm{s,cdc15t}}$ | V | V : 0.97 |
| 94 | *cdc15-ts* multi-copy *TEM1* | $k''_{\mathrm{kp,net}} = 0$, $k_{\mathrm{s,cdc15t}} = 0$,<br>$k_{\mathrm{s,tem1t}} = 5{\times}k_{\mathrm{s,tem1t}}$ | In | V : 0.76 |
| 95 | *cdc15Δ net1-ts* | $k''_{\mathrm{kp,net}} = 0$,<br>$k_{\mathrm{as,rent}}=10$, $k_{\mathrm{as,rentp}}=0.05$,<br>(0.05× WT) | V | V : 0.99 |
| 96 | *cdc15-ts* multi-copy *CDC14* | $k''_{\mathrm{kp,net}} = 0$,<br>$k_{\mathrm{s,14}} = 0.4$ (2 copies) | V | V : 0.96 |
| 97 | *net1-ts* | $k_{\mathrm{as,rent}}=10$, $k_{\mathrm{as,rentp}}=0.05$,<br>(0.05× WT) | V | V : 0.99 |

| | | | | |
|---|---|---|---|---|
| 98 | *GAL-NET1* | $k_{\mathrm{s,net}} = 5\times k_{\mathrm{s,net}}$, $k_{\mathrm{g}} = 0.004621$ | In | In : 0.52 |
| 99 | *cdc14-ts* | $k_{\mathrm{s,14}} = 0$ | In | In : 0.08 |
| 100 | *GAL-CDC14* | $k_{\mathrm{s,14}} = 4\times k_{\mathrm{s,14}}$, $k_{\mathrm{g}} = 0.004621$ | In | In : 0.04 |
| 101 | *GAL-NET1 GAL-CDC14* | $k_{\mathrm{s,net}} = 3\times k_{\mathrm{s,net}}$, $k_{\mathrm{s,14}} = 3\times k_{\mathrm{s,14}}$, $k_{\mathrm{g}} = 0.004621$ | V | V : 0.94 |
| 102 | *net1Δ cdc20-ts* | $k_{\mathrm{as,rent}}$=10, $k_{\mathrm{as,rentp}}$=0.05, (0.05× WT) $k'_{\mathrm{s,20}} = k''_{\mathrm{s,20}} = 0$ | In | In : 0.59 |
| 103 | *cdc14-ts GAL-SIC1* | $k_{\mathrm{s,14}} = 0$, $k'_{\mathrm{s,c1}} = 10\times k'_{\mathrm{s,c1}}$, $k_{\mathrm{g}} = 0.004621$ | In | In : 0.49 |
| 104 | *cdc14-ts* then *GAL-SIC1* | $k_{\mathrm{s,14}} = 0$ for 180 min, then 3 copies of GAL-SIC1 ($k'_{\mathrm{s,c1}} = 3\times k'_{\mathrm{s,c1}}$, $k_{\mathrm{g}} = 0.004621$) | able to exit mitosis | Not able to exit mitosis |
| 105 | *cdc14-ts sic1Δ* | $k'_{\mathrm{s,c1}} = k''_{\mathrm{s,c1}} = 0$, (16% Cdc14 activity) $k_{\mathrm{pp,c1}} = k_{\mathrm{pp,f6}} = 0.64$, $k_{\mathrm{a,swi}} = 0.32$, $k''_{\mathrm{a,cdh}} = 0.128$ | In | In : 0.27 |
| 106 | *cdc14-ts cdh1Δ* | $k_{\mathrm{s,cdh}} = 0$, init CDH1 = CDH1T = 0 (30% Cdc14 activity) $k_{\mathrm{pp,c1}} = k_{\mathrm{pp,f6}} = 1.2$, $k_{\mathrm{a,swi}} = 0.6$, $k''_{\mathrm{a,cdh}}$ =0.13 | In | In : 0.16 |

| | | | | |
|---|---|---|---|---|
| 107 | *cdc14-ts GAL-CLN2* | $k'_{\mathrm{s,n2}} = 0.12$, $k''_{\mathrm{s,n2}} = 0$, $k_{\mathrm{g}} = 0.004621$ (10% Cdc14 activity) $k_{\mathrm{pp,c1}} = k_{\mathrm{pp,f6}} = 0.8$, $k_{\mathrm{a,swi}} = 0.4$, $k''_{\mathrm{a,cdh}} = 0.16$ | In | In : 0.59 |
| 108 | *TAB6*-1 | $k_{\mathrm{as,rent}}$=10, $k_{\mathrm{as,rentp}}$=0.05, (5% WT) | V | V : 0.99 |
| 109 | *TAB6*-1 *cdc15*Δ | $k_{\mathrm{as,rent}}$=10, $k_{\mathrm{as,rentp}}$=0.05, (5% WT), $k''_{\mathrm{kp,net}} = 0$ | V | V : 0.99 |
| 110 | *TAB6*-1 *clb5*Δ *clb6*Δ | $k_{\mathrm{as,rent}}$=10, $k_{\mathrm{as,rentp}}$=0.05, (5% WT), $k'_{\mathrm{s,b5}} = k''_{\mathrm{s,b5}} = 0$ | In | V : 0.99 |
| 111 | *TAB6*-1 *CLB1 clb2*Δ | $k_{\mathrm{as,rent}}$=10, $k_{\mathrm{as,rentp}}$= 0.05, (5% WT), $k'_{\mathrm{s,b2}}$=0.0003, $k''_{\mathrm{s,b2}}$=0.013, (33% WT) | V | V : 0.98 |
| 112 | *mad2*Δ | $k_{\mathrm{mad2}} = 0.01$ | V | V : 0.98 |
| 113 | *bub2*Δ | $k_{\mathrm{bub2}} = 0$, init BUB2 = 0 | V | V : 0.98 |
| 114 | *mad2*Δ *bub2*Δ | $k_{\mathrm{bub2}} = 0$, $k_{\mathrm{mad2}} = 0.01$, init BUB2 = 0 | V | V : 0.99 |
| 115 | *APC-A* | $k''_{\mathrm{a,20}} = 0$ | V | V : 0.97 |
| 116 | *APC-A cdh1*Δ | $k''_{\mathrm{a,20}} = 0$, $k_{\mathrm{s,cdh}} = 0$, init CDH1T = CDH1 = 0 | In | In : 0.18 |
| 117 | *APC-A cdh1*Δ on galactose | $k''_{\mathrm{a,20}} = 0$, $k_{\mathrm{s,cdh}} = 0$, init CDH1T = CDH1 = 0, $k_{\mathrm{g}} = 0.004621$ | In | In : 0.47 |

| | | | | |
|---|---|---|---|---|
| 118 | *APC-A cdh*1Δ multi-copy *SIC*1 | $k''_{\mathrm{a,20}} = 0$, $k_{\mathrm{s,cdh}} = 0$,<br>$k'_{\mathrm{s,c1}} = 15 \times k'_{\mathrm{s,c1}}$,<br>$k''_{\mathrm{s,c1}} = 15 \times k''_{\mathrm{s,c1}}$,<br>init CDH1T = CDH1 = 0 | V | V : 0.83 |
| 119 | *APC-A cdh*1Δ *GAL-SIC*1 | $k''_{\mathrm{a,20}} = 0$, $k_{\mathrm{s,cdh}} = 0$,<br>$k'_{\mathrm{s,c1}} = 10 \times k'_{\mathrm{s,c1}}$,<br>$k_{\mathrm{g}} = 0.004621$,<br>init CDH1T = CDH1 = 0 | V | V : 0.75 |
| 120 | *APC-A cdh*1Δ multi-copy *CDC*6 | $k''_{\mathrm{a,20}} = 0$, $k_{\mathrm{s,cdh}} = 0$,<br>$k'_{\mathrm{s,f6}} = 18 \times k'_{\mathrm{s,f6}}$,<br>$k''_{\mathrm{s,f6}} = 18 \times k''_{\mathrm{s,f6}}$ ,<br>$k'''_{\mathrm{s,f6}} = 18 \times k'''_{\mathrm{s,f6}}$,<br>init CDH1T = CDH1 = 0 | V | V : 0.78 |
| 121 | *APC-A cdh*1Δ *GAL-CDC*6 | $k''_{\mathrm{a,20}} = 0$, $k_{\mathrm{s,cdh}} = 0$,<br>$k'_{\mathrm{s,f6}} = 15 \times k'_{\mathrm{s,f6}}$,<br>$k_{\mathrm{g}} = 0.004621$,<br>init CDH1T = CDH1 = 0 | V | V : 0.82 |
| 122 | *APC-A cdh*1Δ multi-copy *CDC*20 | $k''_{\mathrm{a,20}} = 0$, $k_{\mathrm{s,cdh}} = 0$,<br>$k'_{\mathrm{s,20}} = 25 \times k'_{\mathrm{s,20}}$,<br>$k''_{\mathrm{s,20}} = 25 \times k''_{\mathrm{s,20}}$,<br>init CDH1T = CDH1 = 0 | V | V : 0.79 |
| 123 | *APC-A sic*1Δ | $k''_{\mathrm{a,20}} = 0$, $k'_{\mathrm{s,c1}} = k''_{\mathrm{s,c1}} = 0$ | V | V : 0.83 |
| 124 | *APC-A GAL-CLB*2 | $k''_{\mathrm{a,20}} = 0$, $k'_{\mathrm{s,b2}} = 0.48$,<br>$k_{\mathrm{g}} = 0.004621$ | In | In : 0 |
